# Using child-friendly movie stimuli to study the development of face, place, and object regions from age 3 to 12 years

**DOI:** 10.1101/2021.11.29.469598

**Authors:** Frederik S. Kamps, Hilary Richardson, N. Apurva Ratan Murty, Nancy Kanwisher, Rebecca Saxe

## Abstract

Scanning young children while watching short, engaging, commercially-produced movies has emerged as a promising approach for increasing data retention and quality. Movie stimuli also evoke a richer variety of cognitive processes than traditional experiments – allowing the study of multiple aspects of brain development simultaneously. However, because these stimuli are uncontrolled, it is unclear how effectively distinct profiles of brain activity can be distinguished from the resulting data. Here we develop an approach for identifying multiple distinct subject-specific Regions of Interest (ssROIs) using fMRI data collected during movie-viewing. We focused on the test case of higher-level visual regions selective for faces, scenes, and objects. Adults (N=13) were scanned while viewing a 5.5 minute child-friendly movie, as well as a traditional experiment with isolated faces, scenes, and objects. We found that just 2.7 minutes of movie data could identify subject-specific face, scene, and object regions. While successful, the movie approach was still less effective than a traditional localizer. Having validated our approach in adults, we then used the same methods on movie data collected from 3–12-year-old children (N=122). Movie response timecourses in 3-year-old children’s face, scene, and object regions were already significantly and specifically predicted by timecourses from the corresponding regions in adults. We also found evidence of continued developmental change, particularly in the face-selective posterior superior temporal sulcus. Taken together, our results reveal both early maturity and functional change in face, scene, and object regions, and more broadly highlight the promise of short, child-friendly movies for developmental cognitive neuroscience.

## Introduction

Young children undergo dramatic cognitive development, and a key goal of developmental cognitive neuroscience is to understand the neural correlates of these changes. However, scanning young children with fMRI remains a significant challenge. Consider the literature on the development of higher-level visual cortex, including regions selective for faces, scenes, and objects (Grill-Spector, 2008). Despite a vast behavioral literature suggesting that remarkably mature higher-level visual abilities (e.g., face recognition) emerge well within the first 5 years of life (Nishimura, 2009; McKone et al., 2012), practical limitations have led almost all fMRI studies to date to focus on children 5 years or older (Golarai et al., 2007; Golarai et al., 2010; Cantlon et al., 2011; Scherf et al., 2011; Gomez et al., 2018; Kamps et al., 2020). How much of the picture are we missing without methods for studying early development with fMRI?

One promising approach is to show young children commercially produced movies while scanning with fMRI (Moraczewski, 2018; Cantlon, 2020; Redcay, 2020). This approach trades the rigorous control of traditional experiments for a more engaging (and potentially more ecologically valid) stimulus (Haxby, 2020; Leopold, 2020; Nastase, 2020). While watching movies, young children might be scanned more easily, and provide higher quality data, relative to a traditional design (Vanderwal, 2015). In adults, commercially produced movies can capture remarkably similar functional responses to traditional experimental designs (Hasson et al., 2004; Jacoby, 2016), and the richness of the stimuli enables measuring functional responses during a wide range of cognitive processes simultaneously (Guntupalli, 2016). Applied to developmental populations, naturalistic movies could therefore provide a richer and broader sampling of early functional responses, and thus a greater sensitivity to functional differences between young children and older children or adults (Cantlon, 2013; Moraczewski, 2018; Richardson, 2018; Richardson et al., 2018; Kersey, 2019; Lerner, 2021; Yates, 2021). Indeed, the promise of the naturalistic movie approach has led to initiatives to collect large-scale datasets from children watching movies (Alexander et al., 2017), as well as the development of ever-more-powerful tools for analyzing movie data (Haxby, 2011; Nishimoto S, 2011; Wen, 2018).

Given the increasing supply of fMRI data from young children watching movies, a key next step is to develop and evaluate analysis pipelines that can use these data to make inferences about cognitive and neural function. In particular, our goal was to recover multiple, distinct functional profiles, analogous to those captured in traditional paradigms (e.g., face processing in face regions, or scene processing in scene regions), from a single short movie. Our approach was to define subject-specific regions of interest (ssROIs). ssROIs are among the most simple and powerful tools we have for ensuring that the same functional brain regions are studied across individuals, labs, and experiments, and will therefore be vital for enabling a cumulative research program on the development of specific cortical regions (Saxe et al., 2006; Nieto-Castañón, 2012). Here we therefore ask: can we use short, naturalistic movies suitable for children to define multiple ssROIs with distinct functional profiles? How similar are these functional profiles to those captured by traditional experimental paradigms? And finally, is this approach sufficiently powerful to study the development of functional brain regions in children?

To address these questions, we developed a method for defining ssROIs using just 5.6 minutes of fMRI data collected during movie-viewing and applied it to the test case of higher-level visual regions selective for faces, scenes, and objects. Our approach combines the logic of two previous approaches – intersubject correlations (ISC) (Hasson et al., 2004) and the group-constrained, subject-specific (GSS) ROI definition method (Fedorenko et al., 2010; Julian et al., 2012) – to identify candidate ssROIs based only on that subject’s response to the movie (see Methods). We first validate this movie ssROI approach in adults (N=13), who were scanned while viewing a short, animated, child-friendly movie, as well as a traditional experiment with isolated face, scene, and object conditions, allowing us to directly test the success of the movie ssROI approach. Next, having defined movie ssROIs, we used reverse correlations and deep neural network (DNN) encoding models to assess the extent to which the movie evoked distinct profiles of activation consistent with the well-known functions of these regions. Finally, to explore the power of this approach in data from children directly, we applied the movie ssROI method to an open dataset of children (N=122, ages 3-12 years) who watched the same short movie (Richardson et al., 2018). In so doing, we not only assess the feasibility and sensitivity of our approach in child fMRI data, but also explore the early emergence and development of face, scene, and object regions in higher level visual cortex, which to our knowledge has never been studied with naturalistic movie stimuli, nor with such a large sample or young age group.

## Materials and methods

### Participants and Data

This study involved two datasets; one dataset of adults scanned specifically for this study (N=13; ages 19-35 years; *M*(s.d.) = 26.5(5.3); 6 females), and a second dataset of both children (N=122; ages 3.5-12 years, *M*(s.d.) = 6.7(2.3); 64 females) and adults (N=33; ages 18–39 years; *M*(s.d.) = 24.8(5.3); 20 females), originally collected by Richardson et al., 2018, and freely available from the OpenNeuro database (accession number ds000228). Full details of the subjects and data collection for this dataset can be found in Richardson et al., 2018. All subjects were recruited from the local community, gave written consent (parent/guardian consent and child assent was received for all child subjects), and had normal or corrected to normal vision. Recruitment and experiment protocols were approved by the Committee on the Use of Humans as Experimental Subjects (COUHES) at the Massachusetts Institute of Technology.

### fMRI stimuli

All subjects watched a silent version of ‘Partly Cloudy’ (Reher, 2009), a 5.6-min animated movie. A short description of the plot can be found online (https://www.pixar.com/partly-cloudy#partly-cloudy-1). The stimulus was preceded by 10 s of rest, subtended 17.62 × 13.07 degrees of visual angle.

The adults scanned specifically for this study (N=13), but not those in the Richardson et al. dataset, also completed 2-5 runs of a traditional experiment involving short, dynamic video clips of isolated faces, scenes, objects, and scrambled objects (Kanwisher et al., 1997; Epstein and Kanwisher, 1998; Grill-Spector et al., 1998). For the traditional experiment runs, a blocked design was used in which subjects viewed short videos of faces, objects, scenes, and scrambled objects. Face, object, and scrambled object videos were taken from (Pitcher et al., 2011); scene videos were taken from (Kamps et al., 2016). Each run was 415.8 s long and consisted of 4 blocks per stimulus category. The order of blocks in each run was palindromic (e.g., faces, objects, scenes, scrambled objects, scrambled objects, scenes, objects, faces) and the order of blocks in the first half of the palindromic sequence was pseudorandomized across runs. Each block contained 6, 3-s video clips from the same category, with a 0.3 s interstimulus interval for a total of 19.8 s blocks. Videos subtended 17.76 × 12.89 degrees of visual angle. We also included five 19.8 s fixation blocks: one at the beginning, three in the middle interleaved between each palindrome, and one at the end of each run.

### fMRI data acquisition

Prior to the scan, child subjects completed a mock scan in order to become acclimated to the scanner environment and sounds and to learn how to stay still. Children were given the option to hold a large stuffed animal during the fMRI scan in order to feel calm and to prevent fidgeting. An experimenter stood by child subjects’ feet, near the entrance of the MRI bore, to ensure that the subject remained awake and attentive to the movie. If this experimenter noticed subject movement, she placed her hand gently on the subject’s leg, as a reminder to stay still. Adult subjects were simply instructed to watch the movie and remain still.

For all experiments, whole-brain structural and functional MRI data were acquired on a 3-Tesla Siemens Tim Trio scanner located at the Athinoula A. Martinos Imaging Center at MIT. All subjects used the standard Siemens 32-channel head coil, except for children under age 5, who were scanned using one of two custom 32-channel phased-array head coils made for younger (*N* = 3, *M*(s.d.) = 3.91(.42) years) or older (*N* = 28, *M*(s.d.) = 4.07(.42) years) children (Keil et al., 2011). For child and adult subjects in the Richardson et al. dataset, T1-weighted structural images were collected in 176 interleaved sagittal slices with 1 mm isotropic voxels (GRAPPA parallel imaging, acceleration factor of 3; adult coil: FOV: 256 mm; kid coils: FOV: 192 mm). Functional data were collected with a gradient-echo EPI sequence sensitive to Blood Oxygen Level Dependent (BOLD) contrast in 32 interleaved near-axial slices aligned with the anterior/posterior commissure, and covering the whole brain (EPI factor: 64; TR: 2 s, TE: 30 ms, flip angle: 90**°**). As subjects from this dataset were initially recruited for different studies, there are small differences in voxel size and slice gaps across subjects (3.13 mm isotropic with no slice gap (*N* = 5 adults, *N* = 3 7yos, *N* = 20 8–12yo); 3.13 mm isotropic with 10% slice gap (*N* = 28 adults), 3 mm isotropic with 20% slice gap (*N* = 1 3yo, *N* = 3 4yo, *N* = 2 7yo, *N* = 1 9yo); 3 mm isotropic with 10% slice gap (all remaining subjects)). Prospective acquisition correction was used to adjust the positions of the gradients based on the subject’s head motion one TR back (Thesen, 2000). For adults scanned specifically for this study (n=13), T1-weighted structural images were collected in 176 interleaved sagittal slices with 1 mm isotropic voxels (GRAPPA parallel imaging, acceleration factor of 3; adult coil: FOV: 256 mm (*N* = 8) or 220 mm (*N* = 5)). These functional data were collected with one of two gradient-echo EPI sequences, due to these subjects completing additional scans for an unrelated experiment; one in which 69 slices were oriented approximately between perpendicular and parallel to the calcarine sulcus, covering all of the occipital and temporal lobes, as well as the lower portion of the parietal lobe (*N* = 8; Voxel size: 2×2×2 mm; EPI factor: 64; TR: 2 s, TE: 30 ms, flip angle: 90**°**, slice gap = 0%), or a second with 32 interleaved near-axial slices aligned with the anterior/posterior commissure, and covering the whole brain (*N* = 5; Voxel size: 3×3×3 mm; EPI factor: 84; TR: 2 s, TE: 30 ms, flip angle: 90**°**, slice gap = 10%). For all datasets, functional data were subsequently upsampled in normalized space to 2 mm isotropic voxels. 168 volumes were acquired in each run; children under age five completed two functional runs, while older subjects completed only one run due to time constraints (older participants completed additional, more traditional fMRI experiments). For consistency across subjects, only the first run of data was analyzed. Four dummy scans were collected to allow for steady-state magnetization.

### fMRI data analysis

FMRI data were analyzed using SPM8 (http://www.fil.ion.ucl.ac.uk/spm) (Penny, 2011) and custom software written in Matlab and R. Functional images were registered to the first image of the run; that image was registered to each subject’s anatomical image, and each subject’s anatomical image was normalized to the Montreal Neurological Institute (MNI) template. Previous research has suggested that anatomical differences between children as young as 7 years are small relative to the resolution of fMRI data, which supports usage of a common space between adults and children of this age (for similar procedures with children under age seven, see (Richardson et al., 2018); for methodological considerations, see (Burgund, 2002)). Registration of each individual’s brain to the MNI template was visually inspected, including checking the match of the cortical envelope and internal features like the AC–PC and major sulci. All data were smoothed using a Gaussian filter (5 mm kernel).

Artifact timepoints were identified via the ART toolbox (https://www.nitrc.org/projects/artifact_detect/) (Whitfield-Gabrieli, 2011) as timepoints for which there was (1) >2 mm composite motion relative to the previous timepoint or (2) a fluctuation in global signal that exceeded a threshold of three s.d. from the mean global signal. Subjects were dropped if one-third or more of the timepoints collected were identified as artifact timepoints; this resulted in dropping five child subjects from the original sample (see Richardson et al., 2018). Number of artifact timepoints differed significantly between child and adult subjects in the Richardson et al. dataset (Child (*n* = 122): *M*(s.d.) = 10.5(10.6), Adult (*n* = 33): *M*(s.d.) = 2.8(4), Welch two-sample *t*-test: *t*(137.7) = 6.49, *p* < .000001). Among children, number of motion artifact timepoints was uncorrelated with age (spearman correlation test, (*n* = 122), *r*_s_ (120) = .02, *p* = .86). Notably, mean translation (motion in x, y, and z directions) was similar between adults and children and did not change with age; see Richardson et al., 2018 (Supplemental Figure 8) for visualization of the amount of motion per age group. Despite amount of motion being matched across children, and therefore likely not driving developmental effects within the child sample, we include number of motion artifact timepoints as a covariate in all analyses testing effects of age. Number of artifact timepoints is highly correlated with measures of mean translation, rotation, and distance (*r* > .8). Because this measure is not normally distributed, spearman correlations were used when including amount of motion as a covariate in partial correlations.

All timecourse analyses (including intersubject correlation analyses and reverse correlation analyses) and were conducted by extracting the preprocessed timecourse from each voxel per ROI. We applied nearest neighbor interpolation over artifact timepoints (for methodological considerations on interpolating over artifacts before applying temporal filters, see (Carp, 2013; Hallquist, 2013)), and regressed out two kinds of nuisance covariates to reduce the influence of motion artifacts: (1) motion artifact timepoints; and (2) five principle component analysis (PCA)-based noise regressors generated using CompCor within individual subject white matter masks (Behzadi, 2007). White matter masks were eroded by two voxels in each direction, in order to avoid partial voluming with cortex. CompCor regressors were defined using scrubbed data (e.g., artifact timepoints were identified and interpolated over prior to running CompCor). The residual timecourses were then high-pass filtered with a cutoff of 100 s. Timecourses from all voxels within an ROI were averaged, creating one timecourse per ROI, and artifact timepoints were subsequently excluded (NaNed).

### ROI definition

The study involved three kinds of ROIs: i) group ROIs, ii) traditional ssROIs, and iii) movie ssROIs. All ROIs were initially defined bilaterally, and combined per subject prior to statistical analyses. Group ROIs were defined as all voxels within parcels from a previously published atlas of face (FFA, pSTS), scene (OPA, PPA, RSC) and object (LOC) regions, where each parcel describes the vicinity in which each face, scene, or object ROI is likely to fall given the spatial distribution of individual subject ROIs measured in a large, independent sample of adults (Julian et al., 2012). We chose this particular set of face, scene, and object regions because they are among the best-studied high-level visual regions, and because a reliable group parcel had previously been created for each. A third face region, the occipital face area (OFA), was not included here, since a reliable face-selective response could not be detected in this region. In addition to face, scene, and object regions, group ROIs were also used for theory of mind regions (TPJ, PC, and MPFC) following the methods described in Richardson et al., 2018. A group ROI representing early visual cortex (EVC) was created by drawing a 10mm sphere around peak coordinates generated with Neurosynth (http://neurosynth.org/; [-10 -86 2], [10 -86 2]).

Traditional ssROIs were defined only in the subset of adults (*n*=13) who completed the traditional experiment, using the GSS approach (Fedorenko et al., 2010; Julian et al., 2012). Specifically, for each subject, voxels within each group ROI were ranked based on the t-value of the contrast of either faces > objects (for FFA and pSTS) (Kanwisher et al., 1997), scenes > objects (for OPA, PPA, and RSC) (Epstein and Kanwisher, 1998), objects > scrambled objects (for LOC) (Grill-Spector et al., 1998), or scrambled objects > objects (for early visual cortex, EVC, used to define the movie ssROI, as explained below) (MacEvoy and Yang, 2012; Linsley and MacEvoy, 2014). The traditional ssROI was then defined as the top 100 voxels for the appropriate contrast in each region, following previous approaches (Julian, 2017).

Finally, to define movie ssROIs (in both children and adults), the mean timecourses from traditional ssROIs in adults were regressed on movie data for each subject. For adults who completed the traditional experiment, the mean traditional ssROI timecourses were recalculated separately for each subject after excluding that subject’s own data. Movie ssROIs were then defined using the same GSS procedure as traditional ssROIs, but now with voxels ranked based on contrasts of traditional ROI timecourses. These contrasts were chosen to approximate the contrasts used in a traditional localizer, based on the domain-selectivity of each region. For example, since FFA is defined as faces > objects in a traditional localizer, the movie FFA was defined by the contrast of FFA > LOC. The remaining ROIs were defined using the following contrasts: STS = STS > LOC; OPA = OPA > LOC; PPA = PPA > LOC; RSC = RSC > LOC; LOC = LOC > EVC; EVC = EVC > LOC. We then explored two other approaches to timecourse contrasts. In the first approach, we used contrasts of the mean timecourses of regions averaged by domain; e.g., all face regions > all object regions, all scene regions > all object regions, all object regions > all face and scene regions. In the second approach, we used contrasts of the mean timecourses from each region versus a “counter ROI”, which was defined as the *bottom* 100 voxels (i.e., the least selective voxels) within the search space for each region. Analysis of responses to faces, scenes, objects, and scrambled objects for these approaches revealed at least numerically weaker domain-selectivity, relative to our initial approach, and consequently, these approaches were not considered for further analyses. However, in the process of carrying out the second, “counter ROI” approach, we discovered that including timecourses from the “counter ROIs” in the regression model lead to numerically more selective responses across regions, compared with an analysis that did not include these “counter ROI” timecourses. Consequently, our final regression model for defining movie ssROIs included the mean timecourses from all traditional ssROIs and all “counter ROIs”, and the contrasts used to define movie ssROIs were based on the timecourses from traditional ssROIs only.

### ROI domain-selectivity analyses

To evaluate the success of the movie ROI approach, we compared responses to faces, objects, scenes, and scrambled objects across the three ROI methods using data from the traditional experiment. For each ROI, we extracted beta values (converted to t statistics) for each condition from a standard GLM analysis in SPM. Responses were extracted from each hemisphere separately, and averaged within subjects prior to subsequent analysis.

### Reverse correlation analyses

For reverse correlation analyses, each ROI timecourse in each subject was extracted from each hemisphere, averaged across hemispheres, and z-normalized. Signal values across subjects for each timepoint were tested against baseline (0) using a one-tailed t-test. This procedure is similar to that used by (Hasson et al., 2004). Events were defined as two or more consecutive significantly positive timepoints within each network (i.e., a minimum of 4s in duration). Reverse correlation analysis were conducted in adults (N = 33) and in 3-year-old subjects only (N = 17), in order to examine events that elicit peak responses at maturity and in early childhood. In order to test for developmental effects in the magnitude of response to adult-defined ROI events, we calculated each child’s average response magnitude (from their z-normalized timecourse) across all timepoints identified as an event by each ROI in adults. We then tested for significant correlations between the magnitude of response to adult-defined ROI events and age (as a continuous variable), including amount of motion (number of artifact timepoints) as a covariate. Because this measure of motion is non-normally distributed, we employed spearman correlations.

### Intersubject correlation analyses

In inter-region correlation analysis, the following procedure was used to define ROIs and independently extract timecourses with only a single run of movie data. First, separate movie ssROIs were defined using just the first half (TRs 1-82) or second half of the run (TRs 86-168; the middle 3 time points were not analyzed in order to prevent temporal autocorrelation). Second, the timecourse from the first half of the movie was extracted from the second-half movie ssROIs, and the timecourse from the second half of the movie was extracted from the first-half movie ssROIs (again excluding the middle three time points). Third, the resultant first- and second-half timecourses were concatenated, yielding an estimate of the full movie timecourse. Fourth, timecourses were averaged across hemispheres (e.g., rFFA and lFFA). Fifth, Pearson correlations were calculated between each subject’s movie ssROI timecourse and each mean timecourse from traditional ROIs in adults. Sixth, and finally, the resultant correlations were Fisher z transformed to improve normality for further statistical analysis.

All of the analyses reported in this manuscript should be considered exploratory, not confirmatory, since analyses described here were not chosen prior to data collection, and data collection was not completed with this specific set of analyses in mind. The movie stimulus tested in adults was chosen in order to compare to an existing dataset from children who watched the same movie (rather than based on any particular face, scene, or object content it includes), and we chose analyses based on the stimulus (time series analyses seemed to utilize more data and be more sensitive than previous analysis methods (Jacoby, 2016)), and on recent, relevant progress in the field (Richardson et al., 2018; Yates, 2021).

## Results

### Defining subject-specific ROIs with a short, child-friendly movie

To assess our approach for defining ssROIs using short, child-friendly movie data only, we scanned a group of adults (N=13) while viewing the complete movie once, as well as three runs of a traditional experiment involving dynamic movies of isolated faces, places, and objects. Our analysis focused on two bilateral face regions, the fusiform face area (FFA) and posterior superior temporal sulcus (pSTS); three bilateral scene regions, the occipital place area (OPA), parahippocampal place area (PPA), and retrosplenial complex (RSC); and a bilateral object region, the lateral occipital cortex (LOC). We began by defining face, place, and object regions using data from the traditional experiment (i.e., following standard approaches; see Methods), and extracting response timecourses to the movie from each of these “traditional” ssROIs. We then calculated the average movie timecourse for each region (across N-1 subjects), and regressed these mean “predictor” timecourses on movie data from the left-out subject. Movie ssROIs were defined as the top 100 voxels within a search space showing the highest t-value to the corresponding predictor timecourse (Figure 1).

**Figure 1.**
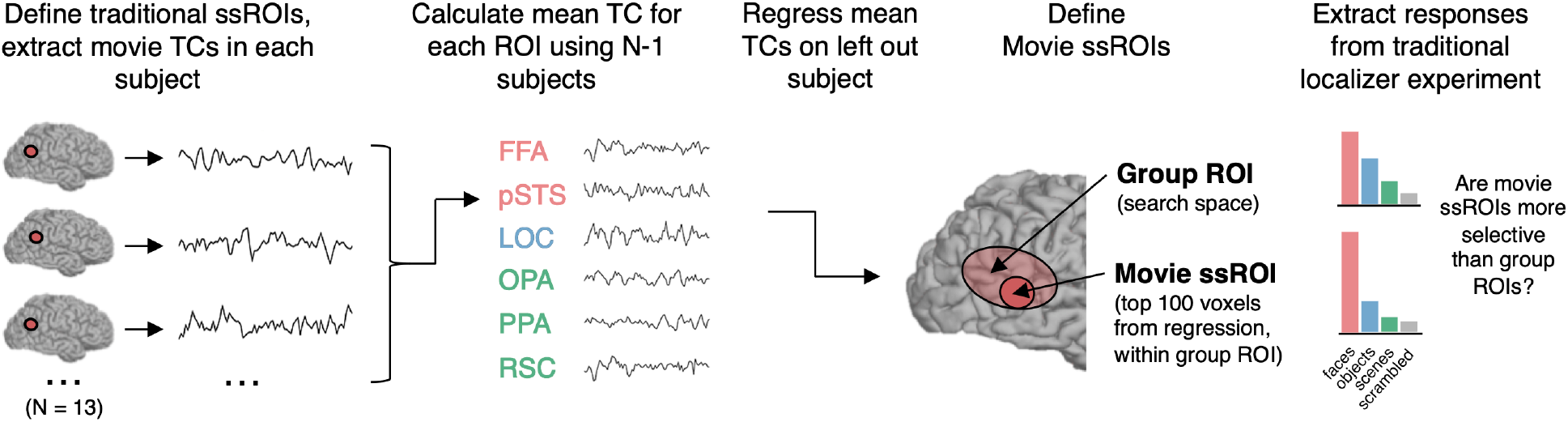
Procedure for defining and evaluating subject-specific functional regions of interest (ssROIs) using only data from a short, child-friendly movie. First, face (i.e., FFA, pSTS), object (i.e., LOC), and scene (i.e., OPA, PPA, RSC) regions were defined in individual adults with traditional stimuli (i.e., “traditional ssROIs”). Second, the timecourse (TC) of response to the child-friendly movie was extracted from each traditional ssROI in each adult, and the mean adult timecourse was calculated using N-1 subjects. Third, the N-1 mean TCs from adults’ traditional ssROIs were regressed per voxel on movie data from the left out subject. Fourth, to define subject-specific movie ROIs (i.e., “movie ssROI”), we selected the top 100 voxels from the regression on that held-out subject (see Methods) within a group search space for each face, scene, or object region. To evaluate whether this procedure successfully defined subject-specific face, scene, and object regions, we extracted responses to faces, scenes, objects, and scrambled objects from the movie ssROIs using data from the traditional experiment, and compared responses to those from group ROIs (i.e., all voxels in the search space for each region, without subject-specific localization), as well as from traditional ssROIs.

A frequently used alternative to ssROIs is to measure activity in ROIs defined by the average activity in an independent group of subjects (“group ROIs”). We asked if the current movie ssROI approach, with limited data, was more effective for identifying the location of functionally selective regions in individual subjects. If the movie ssROI approach is effective, then movie ssROIs will show a stronger difference in response to the conditions that make up the standard localizer contrast for each region (i.e., faces > objects for FFA and pSTS; scenes > objects for OPA, PPA, and RSC; and objects > scrambled objects for LOC), relative to group ROIs. Consistent with this prediction, a 2 (ROI method: movie ssROI vs. group ROI) x 2 (condition) repeated-measures ANOVA revealed a significant ROI method x condition interaction, driven by more domain-selective responses in movie ssROIs than group ROIs, in FFA (*F*_(1,12)_ = 10.32, *p* = 0.007, *η*_*p*_^*2*^= 0.46), OPA (*F*_(1,12)_ = 19.69, *p* < 0.001, *η* _*p*_^*2*^ = 0.62), PPA (*F*_(1,12)_ = 17.74, *p* = 0.001, *η* _*p*_^*2*^ = 0.60), and LOC (*F*_(1,12)_ = 25.72, *p* < 0.001, *η* _*p*_^*2*^ = 0.68). The same analysis revealed marginally significant effects, albeit with large effect sizes, in pSTS (*F*_(1,12)_ = 4.40, *p* = 0.06, *η* _*p*_^*2*^ = 0.27) and RSC (*F*_(1,12)_ = 3.38, *p* = 0.09, *η* _*p*_^*2*^ = 0.22). Full results are shown in Figure 2.

**Figure 2.**
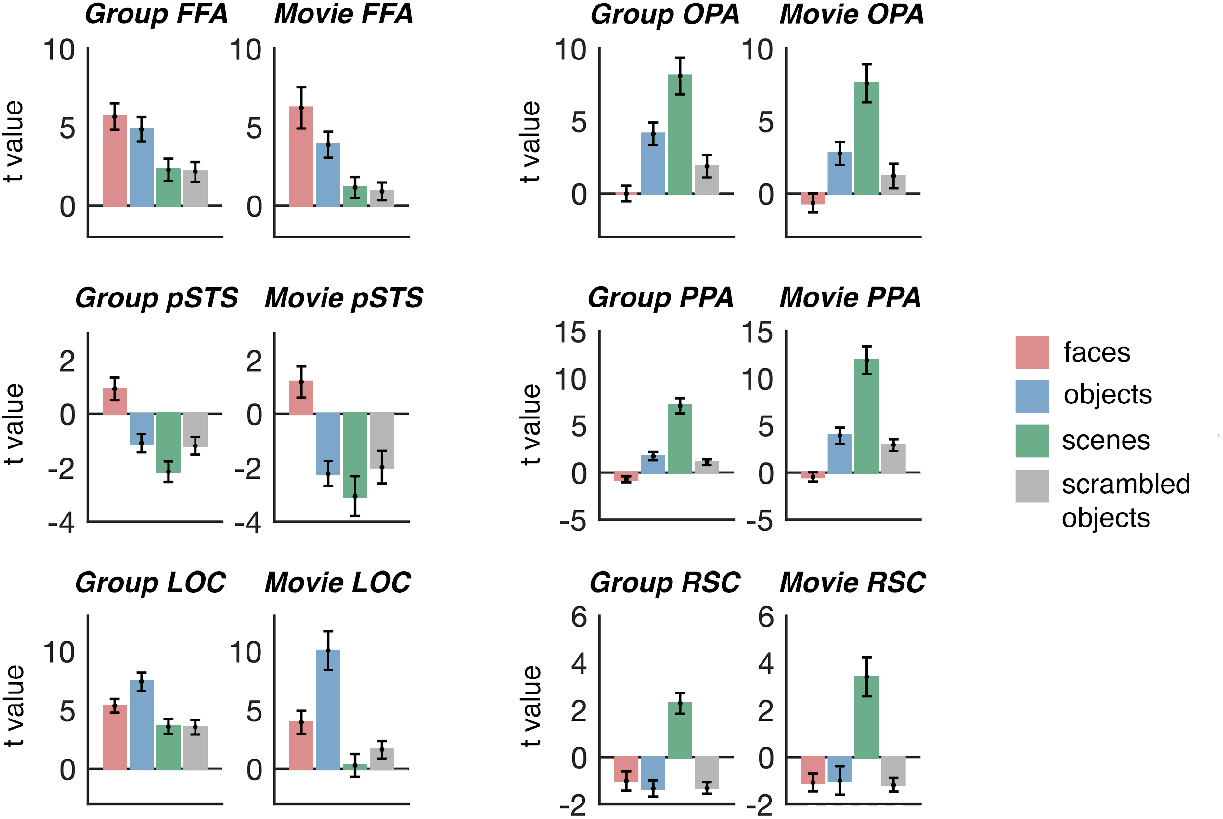
Comparing face, scene, and object selectivity in movie ssROIs versus group ROIs in adults. Each plot shows the response (beta value converted to t statistic) to faces, objects, scenes, and scrambled objects in either group ROIs (left plot for each region) or movie ssROIs (right plot for each region). For most regions, significantly stronger domain-selective responses were observed for movie ssROIs (which involve subject-specific localization) than for group ROIs (which do not involve subject-specific localization).

These results reveal that 5.6 minutes of movie data can be used to identify subject-specific face, scene, and object ROIs. However, to study functional responses in very young children (i.e., <5 years of age), it can be difficult to obtain multiple runs of fMRI data, and especially difficult to obtain data using traditional fMRI experiments. These cases require an approach for defining ROIs based on a limited amount of movie data, such that there is independent data left out in which to further investigate the responses in those ROIs. For this reason, we next asked whether we could use even less movie data – just half of a run, or 2.7 minutes – to localize ROIs. Movie ROIs were defined using the same procedure described above, but now using either the first or second half of the movie only. We then used these movie ssROIs to again extract responses to the four stimulus conditions from the traditional experiment. For statistical analyses, t values for the response to faces, scenes, and objects in the first-half and second-half movie ssROIs were averaged together for each subject, and compared to the responses in group ROIs using the same repeated-measures ANOVA described above. Despite using just 2.7 minutes of movie data, stronger domain-selectivity was measured in movie ssROIs than group ROIs in FFA (*F*_(1,12)_ = 7.242, *p* = 0.019, *η* _*p*_^*2*^ = 0.065), pSTS (*F*_(1,12)_ = 4.929, *p* = 0.046, *η* _*p*_^*2*^ = 0.291), OPA (*F*_(1,12)_ = 13.630, *p* = 0.003, *η* _*p*_^*2*^ = 0.532), PPA (*F*_(1,12)_ = 12.955, *p* = 0.004, *η* _*p*_^*2*^ = 0.519), and LOC (*F*_(1,12)_ = 21.313, *p* < 0.001, *η* _*p*_^*2*^ = 0.640), although no significant effect was observed in RSC (*F*_(1,12)_ = 0.318, *p* = 0.583, *η* _*p*_^*2*^ = 0.026).

How does the domain selectivity of movie ssROIs compare to that for ssROIs defined with a traditional functional localizer for face, scene, and object regions – particularly for cases where a limited amount of data can be collected per subject? To better understand how effectively the movie ssROI approach localizes ssROIs, we next compared adults’ responses to faces, scenes, objects, and scrambled objects in ssROIs defined from the movie versus ssROIs defined using a traditional functional localizer. To approximately match the amount of data used for ROI definition across the two methods, movie ROIs were defined based on the full 5.6-minute movie timecourse, while traditional ROIs were defined using a single 6.3-minute run of the traditional experiment. Response selectivity in each set of ROIs was measured in the remaining runs of the traditional experiment. This analysis revealed significantly more domain-selective responses in the traditional ROIs than the movie ROIs. A 2 (ROI method: movie ssROIs, traditional ssROIs) x 2 (condition) repeated-measures ANOVA found a significant interaction in FFA (*F*_(1,12)_ = 79.329, *p* < 0.001, *η* _*p*_^*2*^ = 0.869), pSTS (*F*_(1,12)_ = 17.884, *p* = 0.001, *η* _*p*_^*2*^= 0.598), OPA (*F*_(1,12)_ = 43.104, *p* < 0.001, *η* _*p*_^*2*^ = 0.872), PPA (*F*_(1,12)_ = 35.622, *p* < 0.001, *η* _*p*_^*2*^ = 0.748), RSC (*F*_(1,12)_ = 53.797, *p* < 0.001, *η* _*p*_^*2*^ = 0.818), and LOC (*F*_(1,12)_ = 54.554, *p* < 0.001, *η* _*p*_^*2*^ = 0.820). Full results are shown in Figure 3. These results underscore the limits of the movie approach: although the current movie localizer is better than a group ROI (i.e., no subject-specific localization whatsoever), traditional localizers with well-controlled, isolated conditions still outperform the current movie approach, even with similar amounts of data.

**Figure 3.**
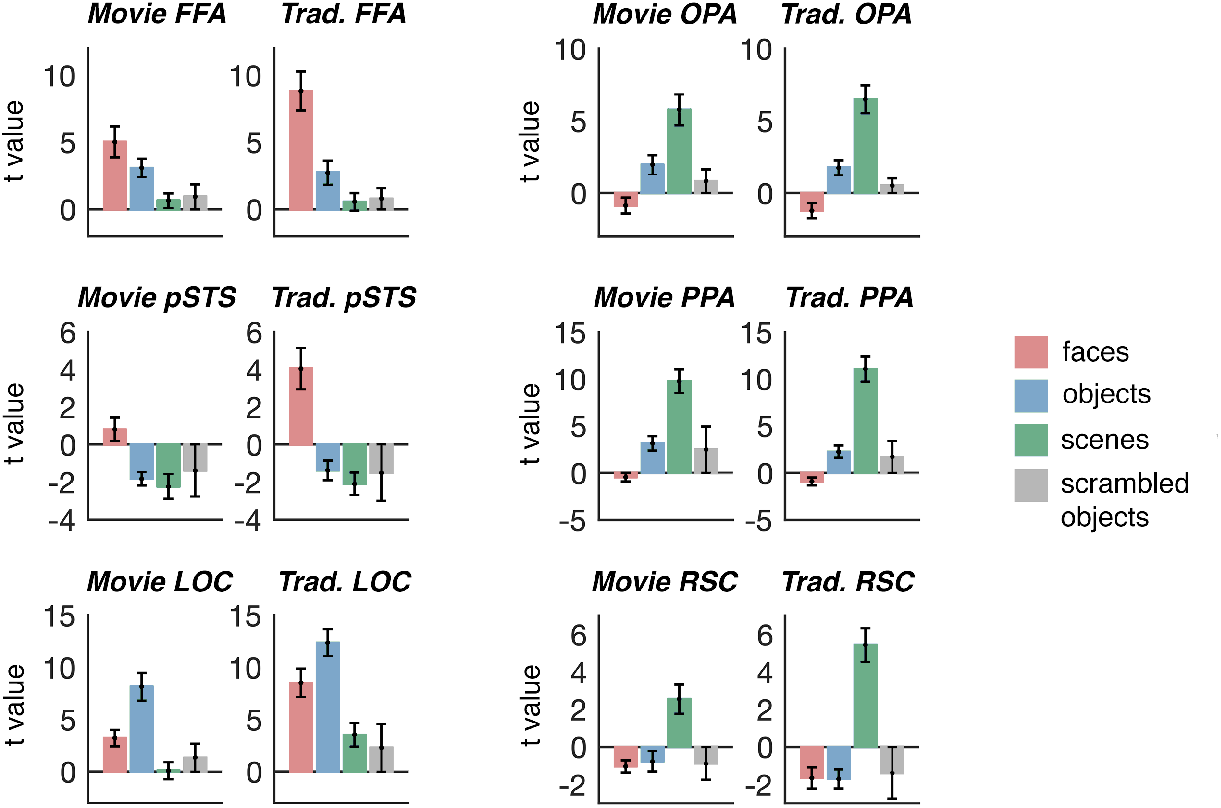
Comparing face, scene, and object selectivity in movie ssROIs versus traditional ssROIs in adults. Each plot shows the response (beta value converted to t statistic) to faces, objects, scenes, and scrambled objects in either movie ROIs (left plot for each region) or traditional ssROIs (right plot for each region). To approximately match the amount of data used for definition of each ROI type, traditional ssROIs were defined using one (6.3 min) run of the traditional experiment (comparable to the 5.6 minute movie), and responses in both ROI types were extracted from the remaining, independent runs of the traditional experiment. For all regions, significantly stronger domain-selective responses were observed for traditional ssROIs than for movie ROIs.

### Exploring functional profiles evoked by a short, child-friendly movie

What content within the movie drives responses within face, scene, and object regions, and how does this content compare with that shown in traditional block designs? If the movie approach is valid, then we should expect this approach to capture similar functions to that captured by traditional approaches; e.g., responses to faces in face areas, responses to scenes in scene areas. At the same time, however, an advantage of child-friendly movies is that they present a broad palate of information that does not neatly fit into four isolated categories. Consequently, child-friendly movies have the potential to “split” related functions that are “lumped” by traditional localizer approaches involving blocked designs with isolated categories (e.g., a movie might reveal dissociations between two face regions, FFA and pSTS, that could not be found in an experiment only testing responses to isolated faces and objects). Addressing these questions, we next performed a series of analyses to validate and explore the functional profiles evoked by the child-friendly movie.

To shed light on what information from the movie stimulus drove responses in each region, we used reversed correlation analysis, a data-driven approach that identifies events (>4s) that elicit strong positive responses in each region. Reverse correlation analysis revealed multiple events for each face, scene, and object region (FFA: 12 events, pSTS: 10 events, OPA: 10 events, PPA: 12 events, RSC: 11 events, LOC: 10 events) (Figure 4). Events largely differed across regions, with greater overlap among regions that share a domain of processing (e.g., two scene regions) than regions that do not (e.g., a face and scene region).

**Figure 4.**
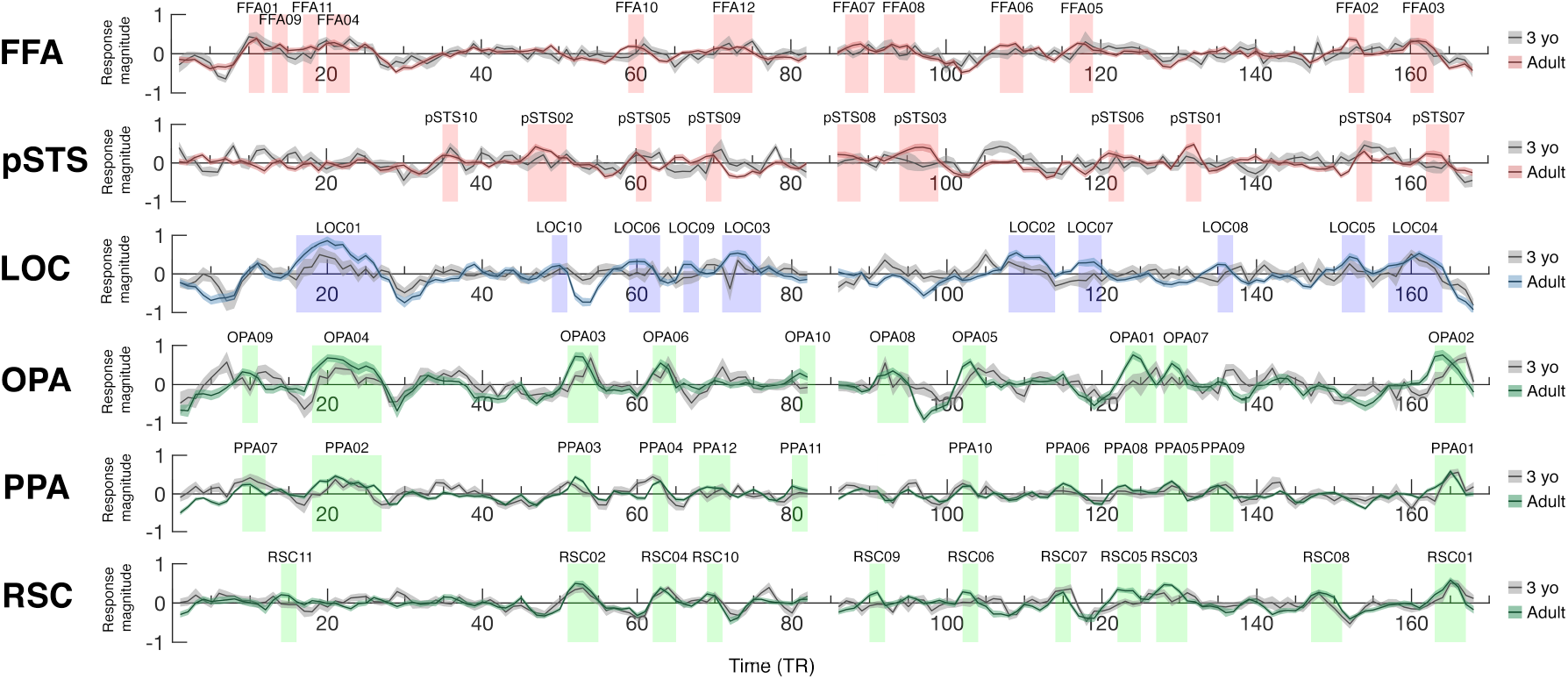
Reverse correlation analysis. The average timecourse from movie ssROIs in adults (colored lines in each plot) and 3 year olds (gray lines in each plot), during viewing of ‘Partly Cloudy’ (Reher, 2009). The dark line indicates the mean response for each group at each time point, and the shaded patch around each line indicates the standard error of the mean. The x-axis depicts the movie time in TRs (2s per TR); the movie experiment was 168 TRs in total (5.6 min). Shaded blocks show time points defined as an event for each region using reverse correlation analysis in adults (N = 33). Event labels (e.g., FFA01, FFA02) indicate ranking of average magnitude of response of peak timepoints in adults.

The content of events appeared to reflect the well-known function of each region, based on visual inspection. For example, FFA events emphasized faces or heads; e.g., the Pixar lamp turns its “head” to look at the viewer; a stork (the main character, “Peck”) knocks the football helmet he is wearing with his fist; Peck and a baby eel look smiling at one another. PPA events emphasized scenes; e.g., the camera zooms out to reveal the world below the clouds; Peck flies alongside a neighborhood of houses; a brief view of the distant scene panning from cloud to cloud. LOC events overlapped strongly with FFA events (not surprisingly, given that timecourses from these regions were highly correlated; see adult-adult intersubject correlations below). However, the order of events eliciting the strongest responses in LOC differed from that in FFA, and events eliciting the strongest LOC responses appeared to involve characters interacting with objects, or handling other animate characters as though they were inanimate objects; e.g., a stork drops off a parcel containing a baby in the window; Peck tosses a baby porcupine from one hand to the other; Gus (a second main character, who is a cloud) rips an alligator off of Peck’s head.

Despite their plausibility, these inferences about what movie content is driving responses are subjective. To provide a more objective way of comparing the functional profiles captured by the movie to that captured in traditional experiments, we used deep convolutional artificial neural network (ANN)-based encoding models of FFA and PPA (Ratan Murty, in press). These encoding models have previously been shown to predict univariate responses in these two regions with remarkable accuracy (models for LOC and other regions tested here have not yet been developed), and reveal striking selectivity for faces and scenes in FFA and PPA, respectively, even across an unprecedented test of selectivity (i.e., across ∼3 million images). Accordingly, these models can offer a precise, quantitative, and unbiased prediction of “expected” face processing in FFA, and scene processing in PPA, even for rich, yet confounded, movie stimuli. Model responses were calculated for each video frame, and averaged across frames in bins of 2s, matching the temporal resolution of the fMRI data (TR = 2s), creating a model “time course”. The FFA encoding model timecourse was significantly positively correlated with the adult FFA (Pearson correlations, *r*(163) = 0.31, *p* < 0.001), but significantly negatively correlated with the adult PPA (*r*(163) = -0.40, *p* < 0.001). By contrast, the PPA encoding model timecourse was significantly positively correlated with the adult PPA (*r*(163) = 0.39, *p* < 0.001), but uncorrelated with the adult FFA (*r*(163) = 0.11, *p* = 0.17). These results provide more objective evidence that response timecourses in FFA and PPA reflect processing of face and scene information, respectively.

Next, to explore whether the movie captured information processing beyond that captured by a traditional experiment with isolated categories, we tested whether the movie captured dissociable responses across the set of face, scene, and object regions, even for regions that share a domain (e.g., FFA and pSTS). To do so, we used an adult-adult intersubject correlation approach. Specifically, we asked if the mean timecourse from traditionally defined ssROIs in one group of adults (N=13; those scanned specifically for this study) showed stronger correlations to that same ROI in a second group of adults’ (N=33, from the Richardson et al. movie dataset) movie-defined ssROIs, compared with another ROI. As can be seen in Figure 6 (“Adult” column, far right), each movie ssROI showed a distinct pattern of correlations across the set of traditional “predictor” timecourses, revealing distinct functional profiles across the regions. The precise pattern in each region reflected dissociations both between regions selective for different domains (e.g., FFA and PPA), and between regions that work on the same domain (e.g., FFA and pSTS). A notable exception was FFA and LOC, which were strongly correlated. This finding may be explained by the fact that very few non-face objects are presented in the movie, as might be required to evoke dissociable responses in these two regions. Nevertheless, a further partial correlation analysis revealed that movie responses even in these highly correlated regions could still be dissociated. For the partial correlation analysis, we recalculated the correlation between timecourses from each traditional ROI and its movie ssROI counterpart after partialling out variance explained by each other traditional ROI. Indeed, all movie ssROIs showed a significant partial correlation with the corresponding traditionally defined ssROI (one-sample t-test, relative to zero; FFA: *t*_(33)_ = 8.30, *p* < 0.001, *d* = 0.25; pSTS: *t*_(33)_ = 11.22, *p* < 0.001, *d* = 0.34; OPA: *t*_(33)_ = 7.85, *p* < 0.001, *d* = 0.24; PPA: *t*_(33)_ = 12.55, *p* < 0.001, *d* = 0.38; RSC: *t*_(33)_ = 10.19, *p* < 0.001, *d* = 0.31; LOC: *t*_(33)_ = 13.42, *p* < 0.001, *d* = 0.41).

### Interregional correlation analyses of movie ROIs in children

The results thus far show that a short, child-friendly movie can be used to define subject-specific face, place, and object regions, and that functional responses to the movie in these regions are not only distinct, but also reflect their well-known functions as captured by traditional paradigms and recent deep convolutional ANN-based encoding models. Moreover, these results establish a means for estimating an entire movie timecourse in independently-defined ssROIs using only a single 5.6-minute run of movie data: the first half of the movie can be extracted from ssROIs defined using the second half of the movie, and the second half of the movie can be extracted from ssROIs defined using the first half (Figure 5). In this final section, we used this approach to explore early emergence and subsequent development of face, scene, and object regions in three- to twelve-year-old children (n=122). In so doing, this analysis allowed us to explore the power of the movie approach for investigating brain development in data-constrained pediatric studies.

**Figure 5.**
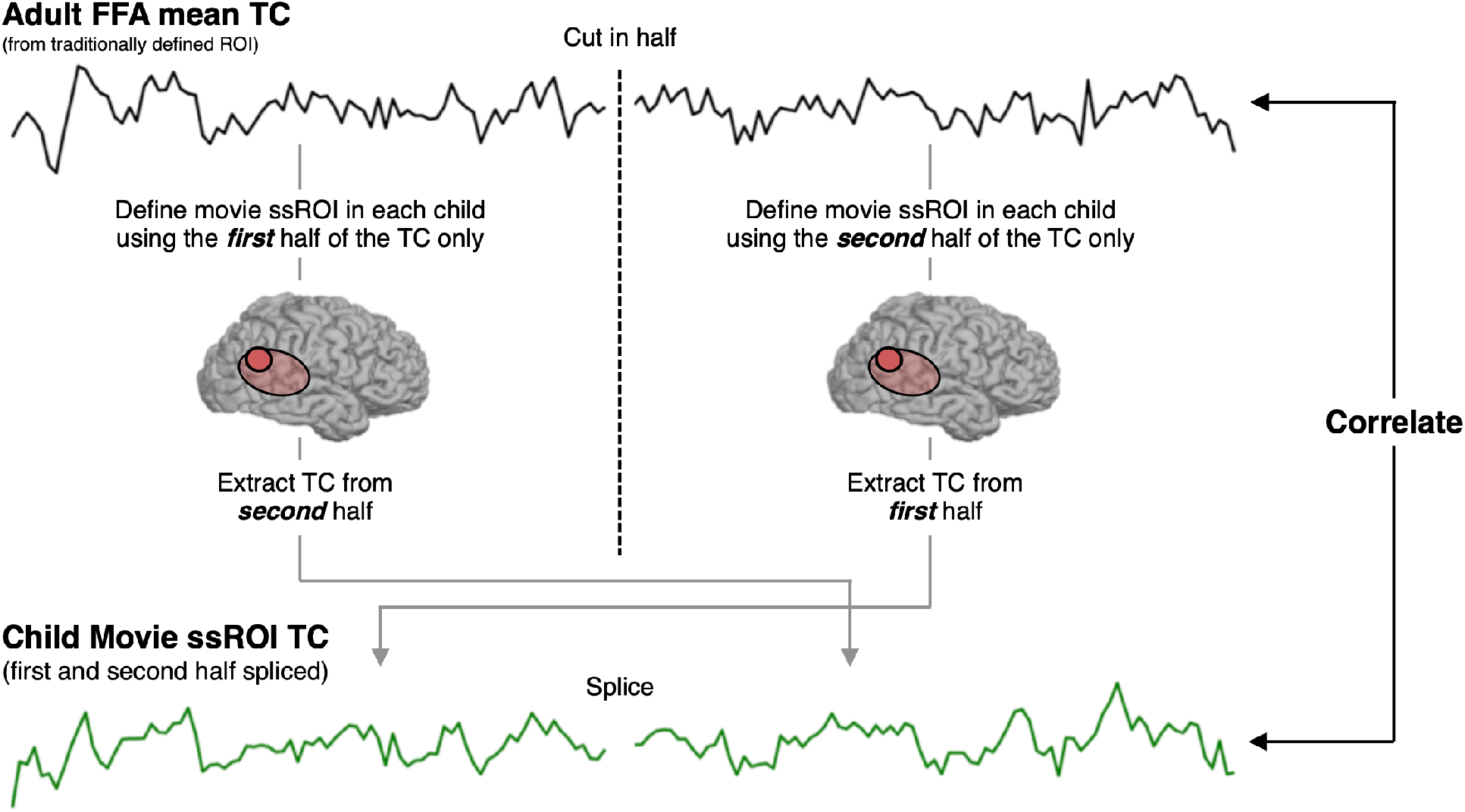
Procedure for child-adult interregional correlation analysis. To investigate whether children show face, scene, and object regions with similar function to adults, we measured the correlation between timecourses from movie ssROIs in children and those from traditionally defined ROIs in adults. Since children only viewed one run of the movie, a split half procedure was used to independently define and test responses. Specifically, the mean adult timecourse from each traditionally defined region (e.g., FFA, as shown here) was split in half (the middle three timepoints were excluded to prevent temporal autocorrelation). The timecourse from the first half of the movie was extracted from movie ssROIs defined using only the second half of the movie, and the timecourse from the second half was extracted from movie ssROIs defined using only the first half. The resultant halves of the movie timecourse were then spliced to form a complete movie ssROI timecourse for each child, and the correlation was measured between these timecourses and those from traditionally defined adult ROIs.

**Figure 6.**
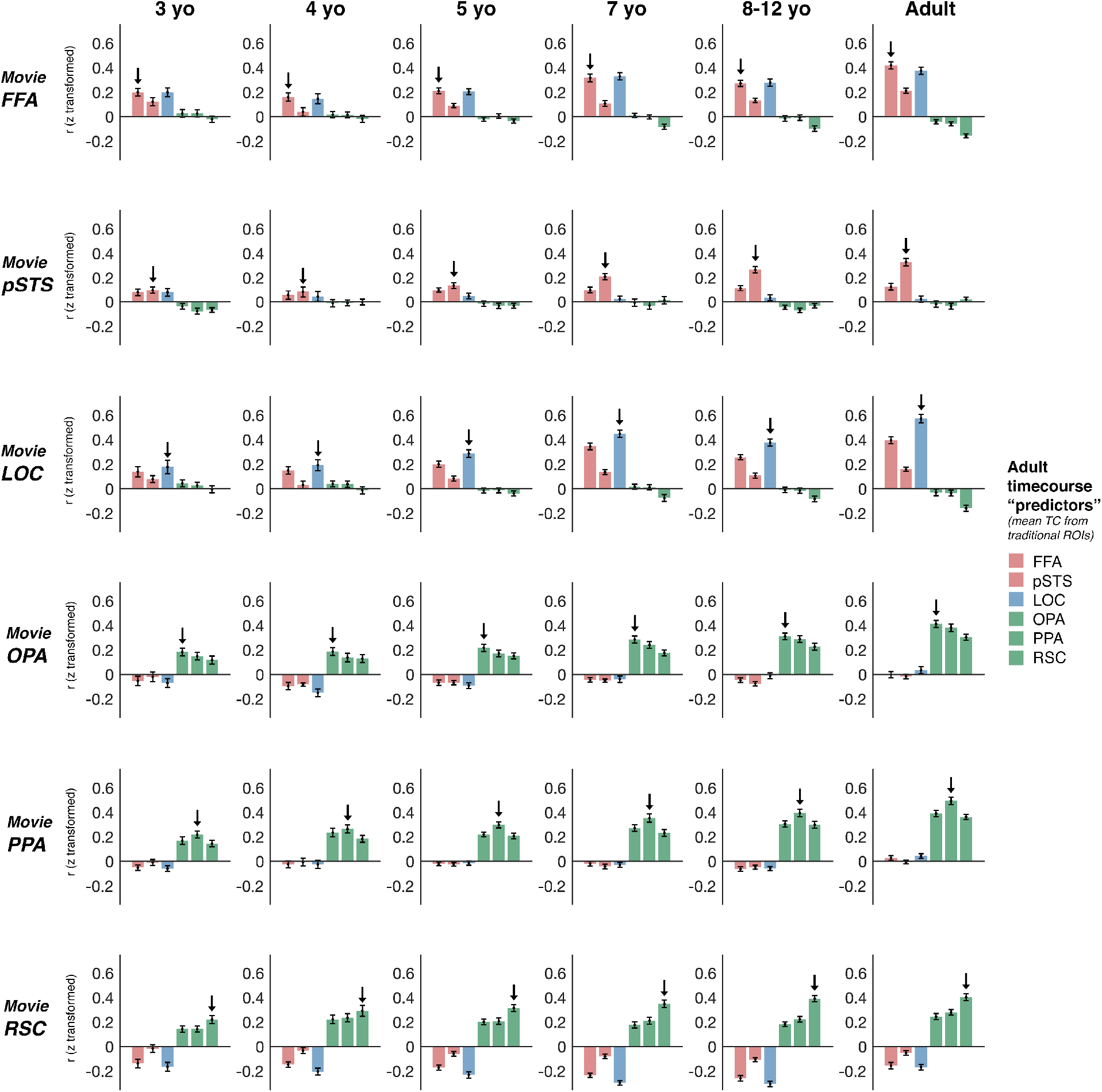
Interregional correlations reveal specific and distinct responses across functional regions, which are early-emerging, but also undergo developmental change throughout childhood. Each plot shows the average correlation of the timecourse from a particular movie ssROI in a particular age group (e.g., the movie FFA in 3 year olds) to the average timecourse from the six traditional ssROIs in adults. Error bars depict the standard error of the mean. Adult movie ssROIs were defined in a different set of adults (*n* = 33) from those in which the average traditional ssROI timecourses were calculated (*n* = 13). For all age groups and regions, the timecourse from movie ssROIs was significantly correlated with the corresponding traditional ssROI timecourse in adults (one-sample t-test, greater than zero; indicated by an arrow on each plot; with the exception of the 4 year old pSTS). These correlations were driven by distinct functional responses to the movie across face, place, and object regions, with movie ssROIs generally better predicted by adult regions with more similar function than adult regions with more distinct function (e.g., scene regions are better predicted by other scene regions than by face regions or object regions). For most regions, a qualitatively similar pattern of results was found in each age group (e.g., for FFA, all age groups show FFA, LOC > pSTS > OPA,PPA,RSC), with the strength of that pattern increasing with age. However, more pronounced developmental change is observed in pSTS, which only showed a greater correlation to the adult pSTS than other regions after age 5.

To test whether children’s movie ssROIs show similar responses to adults, we used child-adult interregional correlations; specifically, we measured the correlation between timecourses extracted from movie ssROIs in children and those from traditional ssROIs in adults (Figure 6). For all regions, timecourses in movie ssROIs were significantly correlated with traditional adult ssROI timecourses by age three years (one-sample t-tests comparing Pearson r values relative to 0; FFA: *t*_(16)_ = 6.34, *p* < 0.001, *d* = 1.54; pSTS: *t*_(16)_ = 3.50, *p* = 0.003, *d* = 0.85; OPA: *t*_(16)_ = 5.81, *p* < 0.001, *d* = 1.41; PPA: *t*_(16)_ = 7.81, *p* < 0.001, *d* = 1.89; RSC: *t*_(16)_ = 6.79, *p* < 0.001, *d* = 1.65; LOC: *t*_(16)_ = 3.17, *p* = 0.006, *d* = 0.77). Similar results were found in all older age groups and ssROIs (all *t*s > 4.61, all *p*s < 0.0001, all *d*s > 1.01; with the exception of a marginal effect in the 4 year old pSTS: *t*_(16)_ = 1.84, *p* = 0.09, *d* = 0.49). The strength of the correlation to the traditional adult timecourse increased with age in all regions (Spearman partial correlation test, including motion (number of artifact timepoints) as a covariate (n=122); FFA: *r*_s_(119) = 0.25, *p* = 0.006; pSTS: *r*_s_(119) = 0.44, *p* < 0.001; OPA: *r*_s_(119) = 0.36, *p* < 0.001; PPA: *r*_s_(119) = 0.35, *p* < 0.001; RSC: *r*_s_(119) = 0.34, *p* < 0.001; LOC: *r*_s_(119) = 0.41, *p* < 0.001).

Importantly, the movie ssROI approach captured distinct functional profiles for each region in children – not just general, non-specific activation to the video (e.g., driven by low-level visual stimulation, or general attention/arousal). This was evident by measuring the correlation of timecourses between each movie ssROI in children and each of the full set of traditional ssROIs in adults. For all regions, and all age groups, a one-way repeated-measures ANOVA revealed a significant main effect of adult ROI (all *F*s > 3.48, all *p*s < 0.013, all *η*_*p*_^*2*^ s > 0.17; with the exception of the 4-year-old pSTS, *F*_(6,96)_ = 1.96, *p* = 0.11, *η* _*p*_^*2*^ = 0.13). As can be seen in Figure 6, each region showed a distinct pattern of correlations to the set of adult regions, and the precise pattern in each region reflected dissociations both between regions that represent the same domain (e.g., FFA and pSTS) and between regions that represent different domains (e.g., FFA and PPA). As in adults, FFA and LOC were strongly correlated at all ages.

Despite the identifiable region-specific responses in children as young as three years, we nevertheless saw clear effects of age; linear mixed effect models testing for effects of adult ROI (i.e., FFA, pSTS, LOC, OPA, PPA, RSC) and age, including motion (number of artifact time points) as a covariate, revealed a significant adult ROI x age interaction in all regions (FFA: *F*_(6,720)_ = 10.61, *p* < 0.001, pSTS: *F*_(6,720)_ = 15.22, *p* < 0.001, OPA: *F*_(6,720)_ = 6.17, *p* < 0.001, PPA: *F*_(6,720)_ = 9.38, *p* < 0.001, RSC: *F*_(6,720)_ = 10.04, *p* < 0.001, LOC: *F*_(6,720)_ = 13.27, *p* < 0.001). In most regions, this age-related change appeared to reflect quantitative increases of otherwise qualitatively similar patterns of correlations in each group (e.g., the movie FFA in every age group showed the same ordering of correlations to adult regions: FFA, LOC > pSTS, EVC > OPA, PPA, RSC). By contrast, in pSTS, the developmental effects included increased differentiation of this region’s functional response. Older children’s pSTS (ages 7 and up) showed a pronounced correlation to the adult pSTS, over and above other regions, whereas younger children’s pSTS (ages 3 and 4) were less functionally distinct, with similar correlations to the adult pSTS, FFA, and LOC.

Can the current movie paradigm reveal even more fine-grained dissociations among these regions in childhood? As in adults, we used a partial correlation approach to provide a stronger test of region-specific function. We recalculated the correlation of timecourses between movie ssROIs in each child and the corresponding, mean traditional ssROI timecourse in adults after partialling out variance explained by the mean timecourse from all other traditional adult ssROI timecourses. In this way, this analysis tests whether children’s movie ssROIs capture the signal unique to each traditional adult ROI. For all regions, 3-year-olds timecourses showed a significant partial correlation with the corresponding region in adults (one-sample t-test, comparing partial Pearson r values relative to 0; pSTS: *t*_(16)_ = 2.14, *p* = 0.049, *d* = 0.13; OPA: *t*_(16)_ = 4.23, *p* < 0.001, *d* = 0.25; PPA: *t*_(16)_ = 8.17, *p* < 0.001, *d* = 0.48; RSC: *t*_(16)_ = 3.62, *p* = 0.002, *d* = 0.21; LOC: *t*_(16)_ = 2.50, *p* = 0.02, *d* = 0.15), with the exception of a marginal effect in FFA (*t*_(16)_ = 2.11, *p* = 0.0502, *d* = 0.12). Similar results were observed in all older groups for all regions (all *t*s > 2.27, all *p*s < 0.05, all *d*s > 0.16), with the exceptions of the 4-year-old pSTS (*t*_(16)_ = 1.33, *p* = 0.21, *d* = 0.10). For most regions, the strength of this region-specific signal increased with age (Spearman partial correlation test, including motion (number of artifact timepoints) as a covariate (n=122)), including pSTS (*r*_s_(119) = 0.45, *p* < 0.001), PPA (*r*_s_(119) = 0.25, *p* = 0.005), RSC (*r*_s_(119) = 0.44, *p* < 0.001), and LOC (*r*_s_(119) = 0.36, *p* < 0.001); however, this analysis did not reveal age-related change in FFA (*r*_s_(119) = -0.03, *p* = 0.71) or OPA (*r*_s_(119) = 0.05, *p* = 0.58).

Does this developmental change reflect quantitative increases of otherwise qualitatively similar movie responses, or do young children’s movie responses differ qualitatively from those in adults? To address this question, we tested for evidence of age-specific function; if young children’s movie responses initially differ reliably from adults, and become increasingly adult-like with age, then we should expect young children to be better predicted by other young children than by adults (and vice versa). To provide the strongest test of these possibilities, we focused on the youngest group of three-year-olds, who should in principle differ most from adults. For each movie ssROI, we calculated the mean timecourse from the three-year-olds and the mean timecourse from movie ssROIs in adults, and measured the correlation of each individual three-year-old’s movie ssROI to these two mean timecourses (each child’s data was always excluded from the calculation of the group mean timecourse). For most regions, three-year-olds’ timecourses were just as strongly correlated to the mean adult timecourse as to the mean three-year-old timecourse (paired samples t-tests, OPA: *t*_(16)_ = 0.30, *p* = 0.76, *d* = 0.07; PPA: *t*_(16)_ = 0.38, *p* = 0.71, *d* = 0.09; RSC: *t*_(16)_ = -0.31, *p* = 0.76, *d* = 0.08; LOC: *t*_(16)_ = -1.83, *p* = 0.09, *d* = 0.44), or more strongly correlated to the adult timecourse than the mean three-year-old timecourse (paired samples t-test, FFA: *t*_(16)_ = -2.90, *p* = 0.01, *d* = 0.70). However, a different pattern was found in pSTS (Figure 8a), where three-year-olds’ timecourses were better predicted by other three-year-olds than by adults (paired samples t-test: *t*_(16)_ = 2.53, *p* = 0.02, *d* = 0.61). To explore how this pSTS result shifted with age, we ran the same analysis, now testing whether each older group (4-, 5-, 7-, and 8–12-year-olds) was more strongly correlated to three-year-olds or to adults. Four-year-olds’ pSTS timecourses were predicted similarly well by 3-year-olds and adults (*t*_(13)_ = -0.34, *p* = 0.74, *d* = 0.09), while 5-year-olds were marginally better predicted by adults (*t*_(13)_ = -1.99, *p* = 0.06, *d* = 0.34), and 7- and 8–12-year-olds were significantly more correlated to adults than 3-year-olds (7-year-olds: *t*_(22)_ = -4.44, *p* < 0.001, *d* = 0.92; 8–12-year-olds: *t*_(33)_ = -8.49, *p* < 0.001, *d* = 1.46). These findings suggest that pSTS undergoes a shift from a reliable early response (present until approximately 3 years) to a reliably different mature response.

To what extent can the developmental trends above be explained by increasing attention to the movie with age? Significant and specific correlations between adult and child timecourses indicate that all groups paid attention to the movie. Nevertheless, it is possible that older participants paid *more* attention than younger participants. This confound can explain general increases (i.e., across all regions) in the ability to detect adult-like function with age, and consequently we do not draw strong conclusions about developmental change in these cases. Critically, however, general attention to the movie cannot fully explain developmental effects in pSTS, specifically, for two reasons. First, increasing attention to the movie with age should affect all regions similarly. By contrast, pSTS showed stronger age-related changes in child-adult intersubject correlations than other regions, including another face region (i.e., FFA; linear mixed effect model testing for effects of region (pSTS, FFA) and age, including motion (number of artifact time points) as a covariate; region x age interaction: *F*_(1,120)_ = 4.37, *p* = 0.03). Second, reduced attention to the movie should lead to weaker or less reliable responses in younger children than older children. However, as reported above, the 3-year-old pSTS was better predicted by other 3 year olds than by adults. In short then, we argue that our study finds clear evidence of developmental change in pSTS, over and above any general, age-related increases in attention.

### Exploring developmental change in pSTS

Having identified a region with reliable developmental differences in functional response to the movie, we explored three (not mutually exclusive) hypotheses about the nature of this developmental change. A first hypothesis is that young children’s’ pSTS initially responds to (at least some) distinct movie events from those in adults, and that responses to movie events eliciting peak responses in adults emerge only gradually with age. We address this possibility in two ways. First, to test whether young children respond to different events from adults, we performed a reverse correlation analysis on data from the youngest children (3 year olds only; timecourses are shown in Figure 4). This analysis identified fewer events than those found in adults in all regions (FFA: 4 events, pSTS: 1 event, OPA: 5 events, PPA: 7 events, RSC: 5 events, LOC: 2 events). For two regions, FFA and LOC, 3-year-old events overlapped entirely with adult events (FFA: 4/4 events, LOC: 2/2 events), emphasizing early emergence and developmental continuity in these regions. For the remaining regions, however, there was at least one 3-year-old event that did not overlap with an adult event (pSTS: 1/1 events, OPA: 2/5 events, PPA: 2/8 events, RSC: 1/5 events). These results provide initial evidence that at least some regions in children – including pSTS and the three scene regions – are driven by different events during the movie than those that drive these regions in adults. However, given that only a small number of such events were identified, future work should seek to replicate this finding, perhaps with larger sample sizes from young children, or longer movies that might capture even more developmental differences. Second, to explore how responses to adult-defined events emerge with age, we measured each child’s mean response magnitude across all adult-defined movie events for each region (Figure 7). In the youngest group of children (3 year olds only), there was a significant positive response to adult-defined movie events in all regions (one-sample t-tests comparing response magnitude relative to zero: FFA: *t*_(16)_ = 4.23, *p* < 0.001, *d* = 1.03; pSTS: *t*_(16)_ = 2.77, *p* = 0.01, *d* = 0.67; OPA: *t*_(16)_ = 4.35, *p* < 0.001, *d* = 1.05; PPA: *t*_(16)_ = 6.43, *p* < 0.001, *d* = 1.56; RSC: *t*_(16)_ = 4.69, *p* < 0.001, *d* = 1.14), with the exception of a marginal effect in LOC (*t*_(16)_ = 2.00, *p* = 0.06, *d* = 0.48). Similar results were found in all older age groups and ssROIs (all *t*s > 2.76, all *p*s < 0.02, all *d*s > 0.74; with the exception of the 4-year-old pSTS: *t*_(13)_ = 1.69, *p* = 0.11, *d* = 0.45). The magnitude of response to the adult-defined movie events increased with age in all regions (Spearman partial correlation test, including motion (number of artifact timepoints) as a covariate (n=122); FFA: *r*_s_(119) = 0.28, *p* = 0.002; pSTS: *r*_s_(119) = 0.51, *p* < 0.001; OPA: *r*_s_(119) = 0.40, *p* < 0.001; PPA: *r*_s_(119) = 0.36, *p* < 0.001; RSC: *r*_s_(119) = 0.39, *p* < 0.001; LOC: *r*_s_(119) = 0.42, *p* < 0.001).

**Figure 7.**
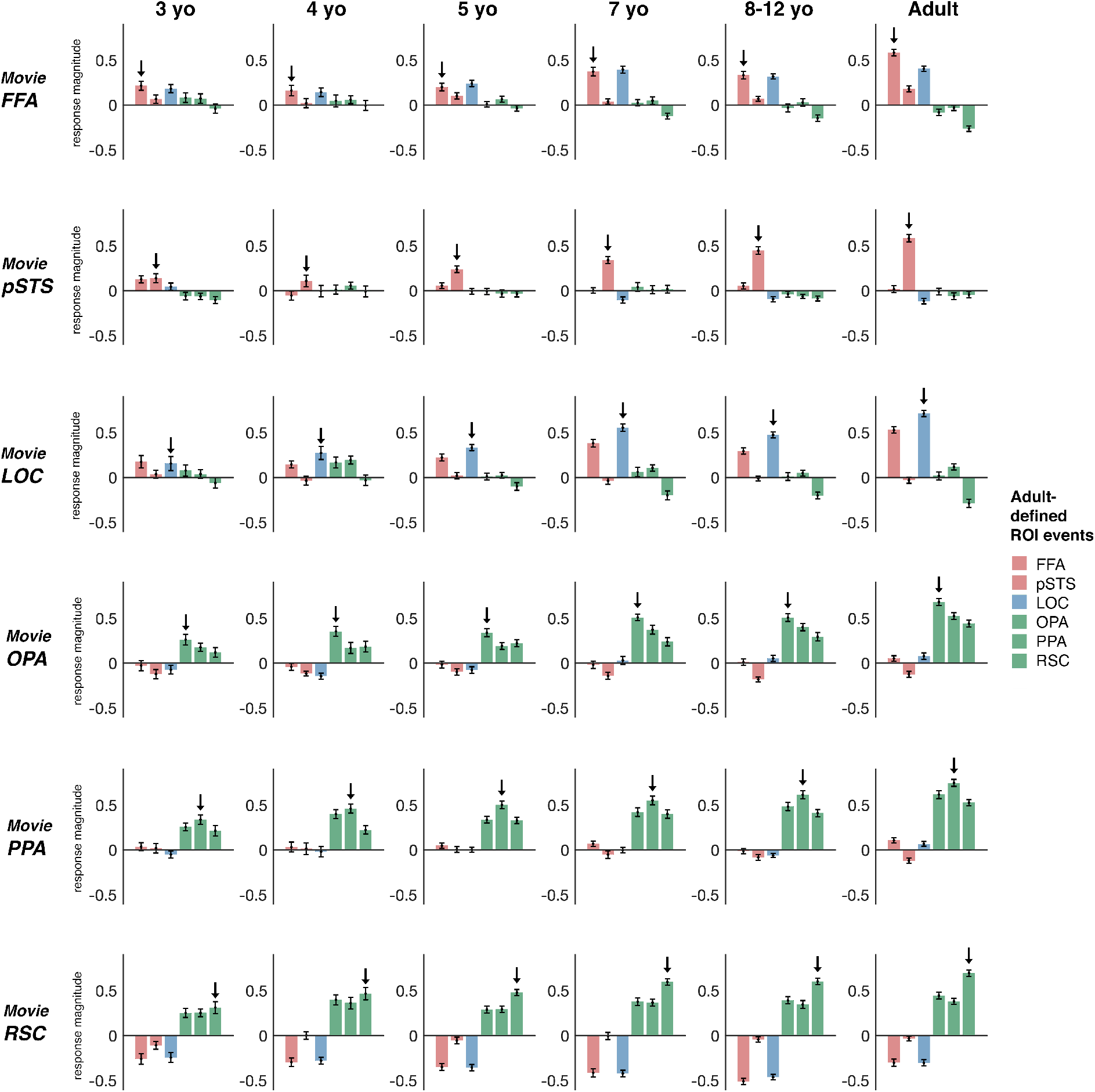
Responses to adult-defined movie events in children. Each plot depicts the mean response of a given movie ssROI in a given age group (e.g., the movie FFA in 3 year olds) to all timepoints defined as an event using reverse correlation analysis in adults. Error bars depict the standard error of the mean. For all age groups, including the youngest group of 3 year olds, movie ssROIs showed significant responses (i.e., greater than 0; indicated by an arrow in each plot) to adult movie events. Responses were generally stronger for events defined by ROIs with more similar functions than ROIs with more distinct functions, mirroring the patterns found in the child-adult interregional correlation analysis. Further, responses to adult events increased significantly with age in most regions, with the clearest effects in pSTS, LOC, and OPA, and with the exception of FFA, which did not show significant age-related change.

**Figure 8.**
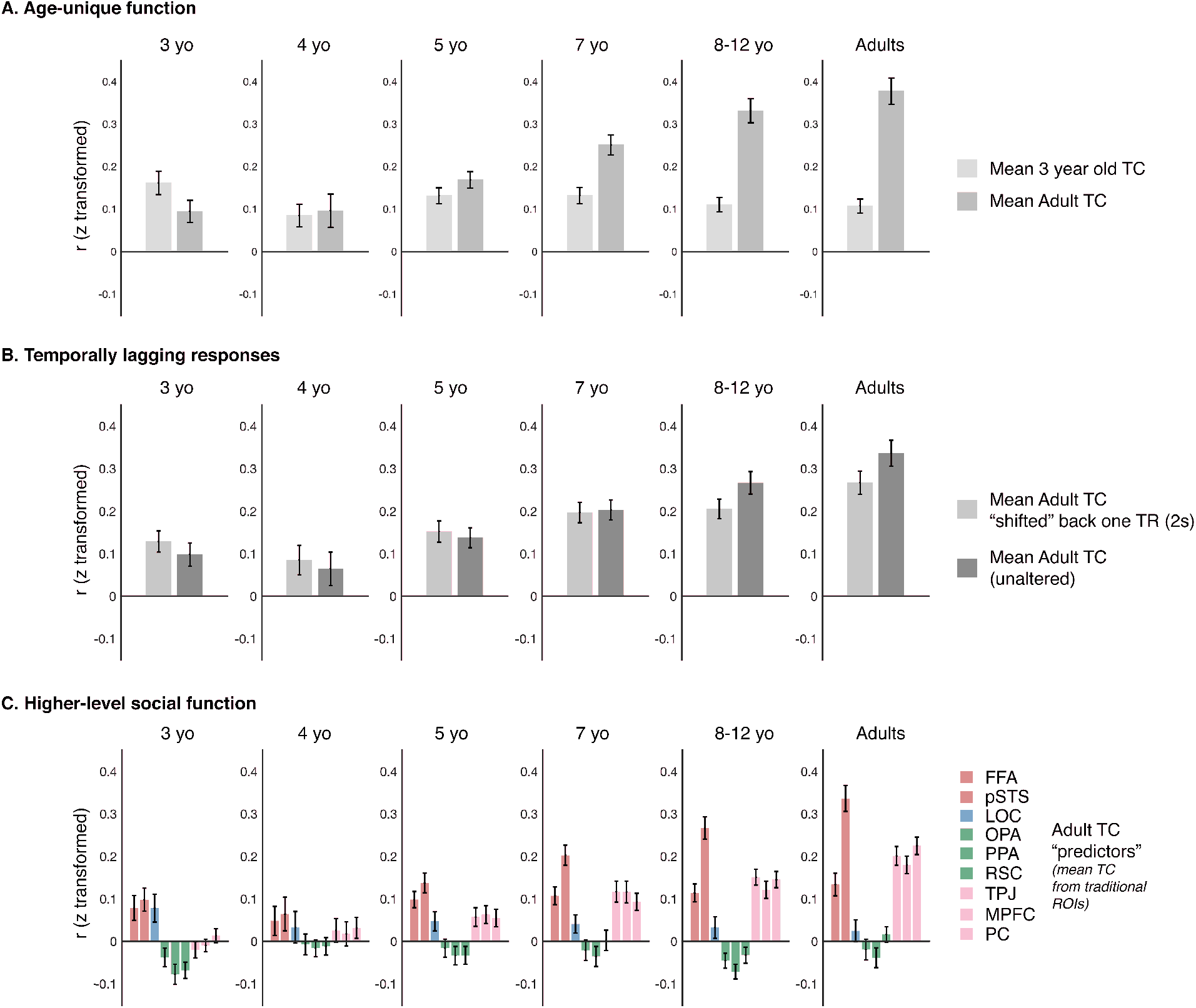
Developmental change in pSTS. The pSTS showed evidence of developmental change across several analyses. (**A**) *Age-unique function*. Each plot depicts the average correlation between timecourses from movie ssROIs in a particular age group and the mean 3-year-old or mean adult timecourse from movie ssROIs. Note that for each 3 year old or adult, the correlation to the mean 3-year-old or adult timecourse was calculated separately for each subject, after excluding that subject from the calculation of the mean timecourse. Movie responses in the 3-year-old pSTS were initially better predicted by the mean 3-year-old than the mean adult timecourse, with the opposite pattern emerging at older ages. (**B**) *Temporally-lagging responses*. Each plot depicts the average correlation between timecourses from movie ssROIs in a particular age group and the mean traditional ssROI timecourse in adults, either temporally shifted backwards by one TR (to simulate a “lagging” response; light orange) or with no temporal shift applied (dark orange). Movie responses in the 3-year-old pSTS are initially better predicted by the shifted mean adult timecourse than by unaltered (non-shifted) timecourse, with the opposite pattern emerging at older ages. (**C**) *Higher-level social function*. Each plot depicts the same child-adult interregional correlations for pSTS shown in Figure 6, but now additionally including mean timecourses from three regions involved in theory of mind processing: the temporal parietal junction (TPJ), Medial Prefrontal Cortex (MPFC), and precuneus (PC). Movie timecourses in the 3-year-old pSTS show stronger correlations to FFA and LOC than to the theory of mind regions. However, this pattern shifts with age, such that older children’s pSTS shows stronger correlations to TPJ, MPFC, and PC than to FFA or LOC. All error bars depict the standard error of the mean.

A second hypothesis is that children’s pSTS responses may lag behind those of adults, because the underlying computational processes, and/or the transmission of information between regions via white matter tracts, are still increasing in efficiency (Wiesmann, 2017). To test this possibility, we compared the correlation of traditional ROI timecourses in adults to those of movie ssROI timecourses in children shifted 1 TR (2s) earlier in time (Figure 8b). For the 3-year-old pSTS, the “shifted” timecourses showed significantly stronger correlations to adult timecourses than non-shifted timecourses (paired samples t-test; *t*_(16)_ = 2.56, *p* = 0.02, *d* = 0.62). Moreover, the strength of this effect diminished continuously with age (linear mixed effect model testing for effects of temporal shift (shifted, not shifted) and age, including motion (number of artifact time points) as a covariate; temporal shift x age interaction: *F*_(1,120)_ = 30.61, *p* < 0.001). By age 8–12, the opposite result was found, with stronger correlations to adult data for the non-shifted timecourse than to the shifted one (paired samples t-test; *t*_(33)_ = -4.54, *p* < 0.001, *d* = 0.78). This effect was specific to pSTS; other regions did not show stronger correlations for shifted than non-shifted timecourses in 3 year olds, and rather showed no effect of the temporal shift, or greater correlations for non-shifted timecourses than shifted ones (as expected given the alignment of timecourses; paired samples t-tests; FFA: *t*_(16)_ = -0.90, *p* = 0.38, *d* = 0.22; PPA: *t*_(16)_ = -5.56, *p* < 0.001, *d* = 1.35; RSC: *t*_(16)_ = -0.96, *p* = 0.35, *d* = 0.23; LOC: *t*_(16)_ = -0.94, *p* = 0.36, *d* = 0.23). One exception was OPA which, like pSTS, showed greater shifted than non-shifted correlations in 3 year olds (paired samples t-test; *t*_(16)_ = 2.61, *p* = 0.02, *d* = 0.63), with the strength of this effect diminishing continuously with age (linear mixed effect model, age by temporal shift interaction: *F*_(1,120)_ = 18.72, *p* < 0.001), such that the 8–12-year-olds showed the opposite effect, albeit marginally (paired samples t-test; *t*_(33)_ = -1.85, *p* = 0.07, *d* = 0.32).

Finally, a third hypothesis is that pSTS takes on increasingly high-level social functions with age. While this hypothesis is challenging to test directly using naturalistic movie data, given that it is difficult to establish precisely what information in the visual display drives responses in a given region (see Discussion), one prediction is that pSTS movie responses will become increasingly similar to those observed in regions recruited for higher-level social cognitive processes (e.g., theory of mind), which also undergo protracted development during childhood (Saxe et al., 2009; Gweon, 2012; Richardson et al., 2018; Richardson, 2020). To test this possibility, we measured the correlation between the pSTS timecourse in children and the mean adult timecourse in three theory of mind regions (bilateral temporal parietal junction, TPJ; precuneus, PC; and medial prefrontal cortex mPFC), defined using group ROIs as reported in Richardson et al. (2018) (Figure 8c). In 3 year olds, the pSTS timecourse was uncorrelated with timecourses of the three adult theory of mind regions (one-sample t-tests comparing Pearson r values relative to 0; TPJ: *t*_(16)_ = -1.01, *p* = 0.33, *d* = 0.25; PC: *t*_(16)_ = -0.64, *p* = 0.53, *d* = 0.16; mPFC: *t*_(16)_ = 0.79, *p* = 0.44, *d* = 0.19). However, by ages 8–12, strong correlations were detectable between pSTS and all three theory of mind regions (one-sample t-tests, relative to 0; TPJ: *t*_(33)_ = 8.06, *p* < 0.001, *d* = 1.38; PC: *t*_(33)_ = 5.95, *p* < 0.001, *d* = 1.02; mPFC: *t*_(33)_ = 7.61, *p* < 0.001, *d* = 1.31), and the strength of these correlations increased significantly with age (spearman partial correlation, including motion artifacts as a covariate; TPJ; *r*_s_(119) = 0.49, *p* < 0.001; PC: *r*_s_(119) = 0.40, *p* < 0.001; mPFC: *r*_s_(119) = 0.43, *p* < 0.001). These effects were specific to pSTS, with no other region showing increasing correlations to theory of mind regions with age (all *r*s < 0.11, all *p*s > 0.24). However, one exception was RSC, which showed increasing correlations to theory of mind regions with age (spearman partial correlation, including motion artifacts as a covariate; TPJ; *r*_s_(119) = 0.44, *p* < 0.001; PC: *r*_s_(119) = 0.38, *p* < 0.001; mPFC: *r*_s_(119) = 0.32, *p* < 0.001), and significant correlations to theory of mind regions in both 8-12 year olds (one-sample t-tests, relative to 0; TPJ: *t*_(16)_ = 13.12, *p* < 0.001, *d* = 2.25; PC: *t*_(16)_ = 10.87, *p* < 0.001, *d* = 1.86; mPFC: *t*_(16)_ = 6.17, *p* < 0.001, *d* = 1.06), and 3 year olds, at least for TPJ and PC (one-sample t-tests, relative to 0; TPJ: *t*_(16)_ = 3.68, *p* = 0.002, *d* = 0.89; PC: *t*_(16)_ = 2.70, *p* = 0.02, *d* = 0.65; mPFC: *t*_(16)_ = -0.23, *p* = 0.82, *d* = 0.05). Critically, it is possible that these effects in RSC may be explained by correlations with the episodic memory network, which is interdigitated with, yet distinct from the ToM network (DiNicola, 2020). Given that the ToM ROIs used here were group defined, rather than functionally defined (as is necessary to distinguish the episodic memory network from the ToM network), our paradigm cannot distinguish these possibilities. Whatever the case for RSC, these analyses support the idea that pSTS development reflects increasingly sophisticated social processing and/or integration with other higher-level social cortical regions with age.

## Discussion

Here we provide an approach for using short, child-friendly movie stimuli to study early brain development with fMRI. We found that even a small amount of movie data can be used to identify subject-specific functional ROIs. Despite the richness of the movie stimuli, we observed distinct activation profiles in face, scene, and object regions that reflected their well-known functions. This method was sufficiently powerful to identify specific regions in children as young as three years old, and also to detect ongoing developmental change, with particularly clear age-related change in the pSTS. Taken together, our results help pave the way for future pediatric studies using movie stimuli, and shed new light on both the early emergence and later development of face, scene, and object regions.

At the broadest level, this work shows that even a single presentation of a 5.6 minute long commercially-produced movie is sufficient for a complete, valid developmental cognitive neuroscience experiment. It is possible to i) define ssROIs, ii) independently extract movie responses from those same ssROIs, iii) explore the nature of the resultant functional profiles, and iv) test the similarity of functional profiles across regions and groups. To facilitate others interested in using this movie ssROI approach, group-average timecourses from all regions tested here are made publicly available with this manuscript. Our success in defining face-, scene-, and object-selective ssROIs complements prior studies showing that this same movie could successfully define ssROIs for regions of the theory of mind network and pain matrix in adults (Jacoby, 2016) and evoke distinct responses in these networks in children (Richardson, 2018; Richardson et al., 2018). Our findings also dovetail more generally with other work harnessing movie data to study predictive processing (Richardson, 2020; Lee, 2021), event structure processing (Baldassano, 2017), and individual differences (Vanderwal, 2017; Richardson, 2018; Finn, 2021). Taken together, these studies highlight the power of even a very short movie to efficiently evoke a broad set of functional profiles, well beyond that possible with a traditional, highly constrained experiment. The potential of the movie approach is further underscored by our findings of reliable differences between young children (i.e., 3 year olds) and adults that might have been missed had we only used a traditional paradigm (i.e., testing a more limited set of conditions with a more constrained set of analyses) or focused only on older children (e.g., > 5 years of age) who are easier to scan.

Our results complement a growing body of work showing dissociable responses among face, place, and object regions by childhood (Golarai et al., 2007; Scherf et al., 2007; Cantlon et al., 2011; Scherf et al., 2011) and infancy (Deen et al., 2017; Powell et al., 2017), but go beyond those studies in two key ways. First, we find such dissociable function using the particularly strong test of a totally uncontrolled naturalistic movie paradigm. That is, despite the lack of constraints for how the stimulus must be viewed (e.g., subjects were not required to maintain fixation), the complex nature of the stimulus (i.e., the movie was not manipulated in any way to better isolate face, scene, or object information), and a difference of more than twenty years visual experience, responses in face, place, and object regions were already strongly correlated with their adult counterparts by three years of age. Second, whereas most previous developmental studies have focused on domain selectivity for faces, scenes, or objects, here we find functional dissociations even between regions that share a domain of information processing (e.g., between FFA and pSTS, which are both selective for faces). This finding suggests that functional dissociations within each network have already begun to emerge by early childhood.

In addition to early emerging function, our study also found evidence of protracted developmental change, particularly in the face-selective pSTS, where developmental change could not be explained by confounds of increasing data quality or attention with age. This finding is consistent with work using traditional paradigms, which has found that responses to faces and socially relevant stimuli are present in pSTS by infancy and early childhood (Otsuka et al., 2007; Lloyd-Fox et al., 2009; Powell et al., 2017; Richardson, 2021), but undergo protracted development late into childhood (Scherf et al., 2007; Ross, 2014; Walbrin, 2020) but see (Golarai et al., 2007). This work also fits with a broader literature suggesting that higher-level social regions in general, beyond the pSTS (e.g., the theory of mind network) undergo protracted development across childhood (Saxe, 2009; Gweon, 2012; Moraczewski, 2018; Richardson et al., 2018; Richardson, 2020). Notably, the movie paradigm used here provides additional insights about the development of pSTS, beyond that captured in previous studies. In particular, we found that the 3-year-old pSTS i) was better predicted by other young children than by adults, ii) showed temporally-lagging responses relative to adults, and iii) showed weaker correlations to other higher-level social regions than adults. These findings support the intriguing possibility that pSTS development involves not only the refinement of an adult-like functional profile, but also more qualitative developmental change. Importantly, however, future work using more tightly controlled paradigms will be required to establish more precisely how the function of pSTS changes across childhood.

Importantly, despite the successes above, this paper also highlights potential shortcomings of the movie approach. For example, although we show that even a short, child-friendly movie can significantly improve ssROI localization, we also show that the movie localizer does not perform as efficiently as a traditional localizer, at least for the particular, short movie we used and the regions we tested here. The advantage of the traditional approach likely stems from the fact that almost every minute is used to measure the contrast-relevant activity, whereas the commercially-produced movie is designed primarily for entertainment. Consequently, we suggest that traditional localizers are still preferable for focused investigations of particular functional regions, especially in populations that can tolerate longer fMRI experiments. An important question for future work will be to investigate whether movie localizers always fall short of traditional localizers, even in typical adults when much more data can be collected per subject. Intriguingly, one study involving ∼10-20x more movie data per subject (i.e., ∼50 to 120 minutes) defined face-selective activation in individual subjects using hyperalignment (Haxby, 2011), and found that that this method identified individual face activation close to or as well as traditional functional localizer experiments (Jiahui, 2020). While promising, however, it is unclear if this result would hold if the amount of movie data and traditional localizer data (i.e., ∼15 to 20 minutes) were matched. In any case, even if traditional localizers ultimately do perform more reliably than movie localizers, there are many contexts in which a short movie localizer might still be preferred; for example, when the experiment requires defining many different ssROIs as efficiently as possible, when working with young children or other populations that better tolerate fMRI experiments, or when analyzing large-scale datasets in which only a single run of a movie, and no additional traditional localizer data, was collected. Thus, researchers will need to consider the details of their particular research question, and the balance of power and flexibility in deciding between these approaches.

Perhaps an even more fundamental limitation of the movie approach, however, lies in the lack of experimental control. Although the reverse correlation and DNN encoding model analyses suggest that movie responses reflect similar functional profiles to those captured in traditional paradigms (e.g., with FFA responding to face information, and PPA responding to scene information), these analyses cannot establish the degree to which responses are driven by i) lower-level information processing (e.g., low-spatial frequency or curvilinear information processing in FFA), ii) domain-selectivity (e.g., selective responses to faces versus scenes and objects), iii) more specific, higher-level functions (e.g., representations of face identity or emotion), or the precise combination thereof. Although one could in theory analyze the movie carefully for each of these kinds of information, and then relate that information to neural responses, the uncontrolled nature of the stimulus would likely always prevent a completely unambiguous inference, particularly with such a short stimulus. Consequently, this work allows for only limited claims about the precise nature of the representations that emerge and change across childhood.

In conclusion, here we explored the promise of naturalistic stimulus approaches for developmental fMRI, focusing on the test case of regions selective for faces, objects, and scenes. We developed and validated an approach for defining subject-specific regions of interest, and then used this approach to define movie ssROIs in children, and explore how responses to a movie stimulus changed over childhood. Remarkably adult-like functional responses were detectable in all regions by just 3 years old, yet these responses also continued to mature across childhood, with the clearest developmental change observed in pSTS.

## Code availability

Summary data necessary to reproduce statistical analyses is available on OSF (https://osf.io/kud5s/). Analysis code used to generate the findings of the study is available from the corresponding author upon request.

## Data availability

The fMRI and behavioral data collected and analyzed during the current study are available through the OpenfMRI project (https://openfmri.org/; Link: https://www.openfmri.org/dataset/ds000228/ DOI: 10.5072/FK2V69GD88). Average movie timecourses from traditionally-defined face, scene, and object ROIs, which can be used to construct regressors for movie localizer experiments in new subjects, are additionally available through OSF (https://osf.io/kud5s/). Additional materials or data are available from the corresponding author upon request.

## Acknowledgments

This work was supported a grant from the Simons Foundation to the Simons Center for the Social Brain (FK), an NSF CAREER award (#095518 to R.S.), an NIH R01-MH096914-05, a Middleton Chair grant (R.S.), and the David and Lucile Packard Foundation (#2008-333024 to R.S.).

## Author Contributions

F.S.K., H.R., and R.S. conceived of and designed the study. Data collection was performed by F.S.K and H.R. Data analysis was performed by F.S.K., H.R., and N.A.R.M., and was supervised by R.S. and N.G.K. The manuscript was drafted by F.S.K., and all other authors provided critical revisions. All authors approved the final version of the manuscript for submission.

## References

Alexander LM et al. (2017) An open resource for transdiagnostic research in pediatric mental health and learning disorders. Sci Data 4.

Baldassano C, Chen, J., Zadbood, A., Pillow, J.W., Hasson, U., Norman, K.A. (2017) Discovering event structure in continuous narrative perception and memory. Neuron 95:709–721.

Behzadi Y, Restom, K., Liau, J. & Liu, T. T. (2007) A component based noise correction method (CompCor) for BOLD and perfusion based fMRI. NeuroImage 37:90–101.

Burgund ED, Kang, H.C., Kelly, J.E., Buckner, R.L., Snyder, A.Z., Petersen, S.E. and Schlaggar, B.L. (2002) The feasibility of a common stereotactic space for children and adults in fMRI studies of development. NeuroImage 1:184–200.

Cantlon JF (2020) The balance of rigor and reality in developmental neuroscience. NeuroImage 216.

Cantlon JF, Pinel P, Dehaene S, Pelphrey KA (2011) Cortical representations of symbols, objects, and faces are pruned back during early childhood. Cerebral cortex 21:191–199.

Cantlon JF, Li, R. (2013) Neural Activity during Natural Viewing of Sesame Street Statistically Predicts Test Scores in Early Childhood. PLoS biology 11:e1001462.

Carp J (2013) Optimizing the order of operations for movement scrubbing: Comment on Power et al. NeuroImage 76:436–438.

Deen B, Richardson H, Dilks DD, Takahashi A, Keil B, Wald LL, Kanwisher N, Saxe R (2017) Organization of high-level visual cortex in human infants. Nature Communications 8.

DiNicola LM, Braga, R. M., & Buckner, R. L. (2020) Parallel distributed networks dissociate episodic and social functions within the individual. Journal of neurophysiology 123:1144–1179.

Epstein, Kanwisher (1998) A cortical representation of the local visual environment. Nature 392:598–601.

Fedorenko E, Hsieh PJ, Nieto-Castanon A, Whitfield-Gabrieli S, Kanwisher N (2010) New method for fMRI investigations of language: defining ROIs functionally in individual subjects. Journal of neurophysiology 104:1177–1194.

Finn ES, Bandettini, P.A. (2021) Movie-watching outperforms rest for functional connectivity-based prediction of behavior. NeuroImage 235:117963.

Golarai G, Liberman A, Yoon JM, Grill-Spector K (2010) Differential development of the ventral visual cortex extends through adolescence. Front Hum Neurosci 3:80.

Golarai G, Ghahremani DG, Whitfield-Gabrieli S, Reiss A, Eberhardt JL, Gabrieli JD, Grill-Spector K (2007) Differential development of high-level visual cortex correlates with category-specific recognition memory. Nature neuroscience 10:512–522.

Gomez J, Natu V, Jeska B, Barnett M, Grill-Spector K (2018) Development differentially sculpts receptive fields across early and high-level human visual cortex. Nature Communications 9.

Grill-Spector K, Kushnir T, Hendler T, Edelman S, Itzchak Y, Malach R (1998) A sequence of object-processing stages revealed by fMRI in the human occipital lobe. Human brain mapping 6:316–328.

Grill-Spector K, Golarai, G., Gabrieli, J. (2008) Developmental neuroimaging of the human ventral visual cortex. Trends in cognitive sciences 12:152–162.

Guntupalli JS, Hanke, M., Halchenko, Y.O., Connolly, A.C., Ramadge, P.J., Haxby, J.V. (2016) A model of representational spaces in human cortex. Cerebral cortex 26:2919–2934.

Gweon H, Dodell-Feder, D., Bedny, M. and Saxe, R. (2012) Theory of mind performance in children correlates with functional specialization of a brain region for thinking about thoughts. Child development 83:1853–1868.

Hallquist MN, Hwang, K. & Luna, B. (2013) The nuisance of nuisance regression: spectral misspecification in a common approach to resting-state fMRI preprocessing reintroduces noise and obscures functional connectivity. NeuroImage 82:208–225.

Hasson U, Nir Y, Levy I, Fuhrmann G, Malach R (2004) Intersubject synchronization of cortical activity during natural vision. Science 303:1634–1640.

Haxby JV, Gobbini, M. I., Nastase, S. A. (2020) Naturalistic stimuli reveal a dominant role for agentic action in visual representation. NeuroImage 216:116561.

Haxby JV, Guntupalli, J.S., Connolly, A.C., Halchenko, Y.O., Conroy, B.R., Gobbini, M.I., Hanke, M., Ramadge, P.J. (2011) A common, high-dimensional model of the representational space in human ventral temporal cortex. Neuron 72:404–416.

Jacoby N, Bruneau, E., Koster-Hale, J., Saxe, R. (2016) Localizing Pain Matrix and Theory of Mind networks with both verbal and non-verbal stimuli. NeuroImage 126:39–48.

Jiahui G, Feilong, M., di Oleggio Castello, M.V., Guntupalli, J.S., Chauhan, V., Haxby, J.V. and Gobbini, M.I. (2020) Predicting individual face-selective topography using naturalistic stimuli. NeuroImage 216.

Julian JB, Fedorenko E, Webster J, Kanwisher N (2012) An algorithmic method for functionally defining regions of interest in the ventral visual pathway. NeuroImage 60:2357–2364.

Julian JB, Ryan, J., Epstein, R.A. (2017) Coding of Object Size and Object Category in Human Visual Cortex. Cerebral cortex 27:3095–3109.

Kamps, Lall V, Dilks DD (2016) Occipital place area represents first-person perspective motion through scenes. Cortex; a journal devoted to the study of the nervous system and behavior 83:17–26.

Kamps, Pincus JE, Radwan SF, Wahab S, Dilks DD (2020) Late Development of Navigationally Relevant Motion Processing in the Occipital Place Area. Curr Biol.

Kanwisher N, McDermott J, Chun MM (1997) The fusiform face area: a module in human extrastriate cortex specialized for face perception. J Neurosci 17:4302–4311.

Keil B, Alagappan V, Mareyam A, McNab JA, Fujimoto K, Tountcheva V, Triantafyllou C, Dilks DD, Kanwisher N, Lin W, Grant PE, Wald LL (2011) Size-optimized 32-channel brain arrays for 3 T pediatric imaging. Magn Reson Med 66:1777–1787.

Kersey AJ, Wakim, K.M., Li, R. and Cantlon, J.F. (2019) Developing, mature, and unique functions of the child’s brain in reading and mathematics. Developmental Cognitive Neuroscience 39:100684.

Lee CS, Aly, M., Baldassano, C. (2021) Anticipation of temporally structured events in the brain. Elife 10:e64972.

Leopold DAaP, S.H. (2020) Studying the visual brain in its natural rhythm. NeuroImage 216:116790.

Lerner Y, Scherf, S. K., Katkov, M., Hasson, U., Behrmann, M. (2021) Changes in Cortical Coherence Supporting Complex Visual and Social Processing in Adolescence. J Cognitive Neurosci:1–16.

Linsley D, MacEvoy SP (2014) Evidence for participation by object-selective visual cortex in scene category judgments. :Journal of vision 14.

Lloyd-Fox S, Blasi A, Volein A, Everdell N, Elwell CE, Johnson MH (2009) Social perception in infancy: a near infrared spectroscopy study. Child development 80:986–999.

MacEvoy SP, Yang Z (2012) Joint neuronal tuning for object form and position in the human lateral occipital complex. NeuroImage 63:1901–1908.

McKone E, Crookes K, Jeffery L, Dilks DD (2012) A critical review of the development of face recognition: Experience is less important than previously believed. Cogn Neuropsychol 29:174–212.

Moraczewski D, Chen, G., Redcay, E. (2018) Inter-subject synchrony as an index of functional specialization in early childhood. Scientific Reports 8:1–12.

Nastase SA, Goldstein, A., Hasson, U. (2020) Keep it real: rethinking the primacy of experimental control in cognitive neuroscience. NeuroImage 222:117254.

Nieto-Castañón A, & Fedorenko, E. (2012) Subject-specific functional localizers increase sensitivity and functional resolution of multi-subject analyses. NeuroImage 63:1646–1669.

Nishimoto S VA, Naselaris T, Benjamini Y, Yu B, Gallant JL. 2 (2011) Reconstructing visual experiences from brain activity evoked by natural movies. Curr Biol 21:1641–1646.

Nishimura M, Scherf, S. and Behrmann, M. (2009) Development of object recognition in humans. :F1000 biology reports 1.

Otsuka Y, Nakato E, Kanazawa S, Yamaguchi MK, Watanabe S, Kakigi R (2007) Neural activation to upright and inverted faces in infants measured by near infrared spectroscopy. NeuroImage 34:399–406.

Penny WD, Friston, K.J., Ashburner, J.T., Kiebel, S.J. and Nichols, T.E. eds. (2011) Statistical parametric mapping: the analysis of functional brain images. Elsevier.

Pitcher D, Dilks DD, Saxe RR, Triantafyllou C, Kanwisher N (2011) Differential selectivity for dynamic versus static information in face-selective cortical regions. NeuroImage 56:2356–2363.

Powell LJ, Deen B, Saxe R (2017) Using individual functional channels of interest to study cortical development with fNIRS. Developmental science 21:e12595.

Ratan Murty NA, Bashivan, P., Abate, A., DiCarlo, J. J., Kanwisher, N. (in press) Computational models of category-selective brain regions enable high-throughput tests of seletivity. :Nat Commun.

Redcay E, Moraczewski, D. (2020) Social cognition in context: A naturalistic imaging approach. NeuroImage 216:116392.

Reher K, & Sohn, P. (2009) Partly Cloudy. In.

Richardson H (2018) Development of brain networks for social functions: Confirmatory analyses in a large open source dataset. :Dev Cogn Neurosci.

Richardson H, Lisandrelli G, Riobueno-Naylor A, Saxe R (2018) Development of the social brain from age three to twelve years. :Nature Communications 9.

Richardson H, Taylor, J., Kane-Grade, F., Powell, L., Bosquet Enlow, M., Nelson, C.A. (2021) Preferential responses to faces in superior temporal and medial prefrontal cortex in three-year-old children. Developmental Cognitive Neuroscience 50:100984.

Richardson HaS, R. (2020) Development of predictive responses in theory of mind brain regions. Developmental science 23:e12863.

Ross PD, de Gelder, B., Crabbe, F. and Grosbras, M.H. (2014) Body-selective areas in the visual cortex are less active in children than in adults. Front Hum Neurosci 8:941.

Saxe R, Brett M, Kanwisher N (2006) Divide and conquer: a defense of functional localizers. NeuroImage 30:1088–1096; discussion 1097-1089.

Saxe RR, Whitfield-Gabrieli S, Scholz J, Pelphrey KA (2009) Brain regions for perceiving and reasoning about other people in school-aged children. Child development 80:1197–1209.

Saxe RR, Whitfield-Gabrieli, S., Scholz, J. and Pelphrey, K.A. (2009) Brain regions for perceiving and reasoning about other people in school-aged children. Child development 80:1197–1209.

Scherf KS, Behrmann M, Humphreys K, Luna B (2007) Visual category-selectivity for faces, places and objects emerges along different developmental trajectories. Developmental science 10:F15–30.

Scherf KS, Luna B, Avidan G, Behrmann M (2011) “What” Precedes “Which”: Developmental Neural Tuning in Face-and Place-Related Cortex. Cerebral cortex 21:1963–1980.

Thesen S, Heid, O., Mueller, E. & Schad, L. R. (2000) Prospective acquisition correction for head motion with image-based tracking for real-time fMRI. Magn Reson Med 44:457–465.

Vanderwal T, Eilbott, J., Finn, E., Craddock, R.C., Turnbull, A., Castellanos, F.X. (2017) Individual differences in functional connectivity during naturalistic viewing conditions. NeuroImage 157:521–530.

Vanderwal T, Kelly, C., Eilbott, J., Mayes, L. C., & Castellanos, F. X. (2015) Inscapes: A movie paradigm to improve compliance in functional magnetic resonance imaging.. NeuroImage 122:222–232.

Walbrin J, Mihai, I., Landsiedel, J., Koldewyn, K. (2020) Developmental changes in visual responses to social interactions. Developmental Cognitive Neuroscience 42:100774.

Wen H, Shi, J., Zhang, Y., Lu, K.H., Cao, J., Liu, Z. (2018) Neural Encoding and Decoding with Deep Learning for Dynamic Natural Vision. Cerebral cortex 28:4136–4160.

Whitfield-Gabrieli S, Nieto-Castanon, A. and Ghosh, S. (2011) Artifact detection tools (ART). In. Cambridge, MA.

Wiesmann CG, Schreiber, J., Singer, T., Steinbeis, N. & Friederici, A.D. (2017) White matter maturation is associated with the emergence of Theory of Mind in early childhood. :Nat Commun 8.

Yates TS, Ellis, C.T. and Turk-Browne, N.B. (2021) Emergence and organization of adult brain function throughout child development. NeuroImage 226:117606.

